# Mechanistic insights into dideoxygenation in gentamicin biosynthesis

**DOI:** 10.1101/2021.05.12.443773

**Authors:** Sicong Li, Priscila dos Santos Bury, Fanglu Huang, Junhong Guo, Guo Sun, Anna Reva, Chuan Huang, Xinyun Jian, Yuan Li, Jiahai Zhou, Zixin Deng, Finian J. Leeper, Peter F. Leadlay, Marcio V. B. Dias, Yuhui Sun

**Author notes:** These authors contributed equally (Sicong Li, Priscila dos Santos Bury, Fanglu Huang).

## Abstract

Gentamicin is an important aminoglycoside antibiotic used for treatment of infections caused by Gram-negative bacteria. Although most of the biosynthetic pathway of gentamicin has been elucidated, a remaining intriguing question is how the intermediates JI-20A and JI-20B undergo a dideoxygenation to form gentamicin C complex. Here we show that the dideoxygenation process starts with GenP-catalyzed phosphorylation of JI-20A and JI-20Ba. The phosphorylated products are converted to C1a and C2a by concerted actions of two PLP (pyridoxal 5’-phosphate)-dependent enzymes: elimination of water and then phosphate by GenB3 and double bond migration by GenB4. Each of these reactions liberates an imine which hydrolyses to a ketone or aldehyde and is then re-aminated by GenB3 using an amino donor. Crystal structures of GenB3 and GenB4 have guided site-directed mutagenesis to reveal crucial residues for the enzymes’ functions. We propose catalytic mechanisms for GenB3 and GenB4, which shed new light on the already unrivalled catalytic versatility of PLP-dependent enzymes.

---

Aminoglycosides are antibiotics that kill most Gram-negative bacteria by interfering with protein synthesis^1,2,3^. Gentamicin is a member of the 2-deoxystreptamine (2-DOS)-containing aminoglycoside family. Clinically used gentamicin C is a mixture containing five compounds: C1, C1a, C2, C2a, and C2b, which share a structure consisting of a 2-DOS aglycone (ring I) with a purpurosamine (ring II) and garosamine (ring III) attached at C-4 and C-6, respectively, but which differ from each other in the methylations on ring II. All five components lack the C-3’ and C-4’ hydroxyl groups (Fig. 1), which protects them from being inactivated by some aminoglycoside modifying enzymes (AMEs)^4,5,6,7^. Since the biosynthetic gene cluster of gentamicin was identified more than a decade ago, we and others have elucidated most of the gentamicin biosynthetic pathway in *Micromonospora echinospora* ATCC15835^8–19^ and revealed crystal structures of several of the enzymes involved in the biosynthesis^14,20,21,22^. GenD2, GenN, GenS2 and GenD1 have been demonstrated to be involved in the formation of gentamicin X2^16,21,22^. At X2 there is a branching of the pathway: methylation of X2 at C-6’ by GenK produces G418, precursor of gentamicins C1, C2 and C2a,^13,14,16^ whereas this methylation does not occur on route to gentamicins C1a and C2b (Fig. 1). X2 and G418 undergo oxidation followed by amination at C-6’ by GenQ and GenB1 to generate JI-20A (**1**) and JI-20Ba (**2**), respectively^15^. GenB2 exhibits 6’-epimerase activity primarily interconverting C2a and C2^15^, but able to also epimerise JI-20Ba (**2**) to JI-20Bb^19^. The N-6’ methyl groups of C2b and C1 are added by GenL, a methyltransferase unusually located 2.54 Mbp away from the known gentamicin gene cluster^17^. However, the enzymes responsible for the 3’,4’ didehydroxylation of **1** and **2** leading to C1a and C2a are still enigmatic, although GenP, GenB3 and GenB4 have been suggested as the likely candidates^15,18,19,23^. Particularly, GenB3 and GenB4 belong to the PLP-dependent enzyme family, which has been reported to catalyze various reactions, such as transamination, isomerization, desulfurization and decarboxylation^24^.

**Fig. 1.**
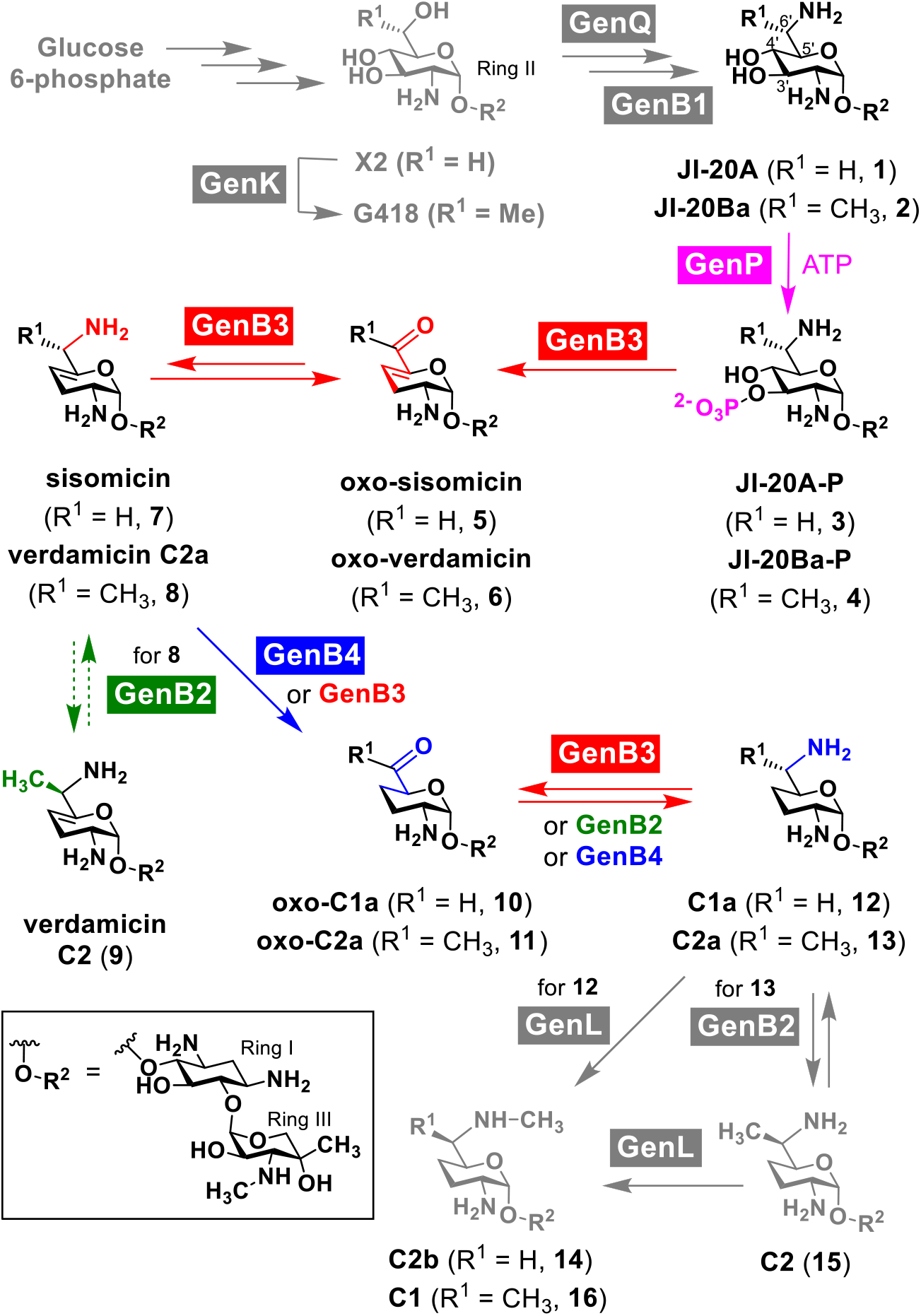
The biosynthetic pathway for the 3’,4’-didehydroxylation and 4’,5’-double bond reduction in gentamicin biosynthesis. The structures and the corresponding enzymes, GenP, GenB2, GenB3 and GenB4, related to this study are indicated in pink, green, red and blue, respectively. Dashed arrows refer to conversion without substantial experimental evidence. The pathway presented in grey has been reported previously.

In the present study, in-frame gene deletion, *in vivo* feeding of intermediates, *in vitro* enzyme assays and protein crystallography confirm that GenP, a homologue of aminoglycoside 3’-phosphotransferases, and two PLP-containing enzymes, GenB3 and GenB4, are responsible for the conversion of **1** and **2** to C1a (**12**) and C2a (**13**), respectively. Catalytic mechanisms for GenB3 and GenB4 are also proposed.

## Results

### 3’,4’-didehydroxylation of 1 and 2 is catalyzed by successive actions of GenP and GenB3

The accumulation of **1** and **2** observed in a *genP* deletion mutant^19^ and the ability of a recombinant GenP to phosphorylate *in vitro* the 3’-OH of kanamycin B^23^, an analogue of **1** and **2**, has led to speculation that phosphorylation of **1** and **2** by GenP is a prerequisite for their 3’,4’-didehydroxylation. To verify the hypothesis, several in-frame deletion mutants were constructed and their culture extracts were analyzed by LC-ESI-HRMS. Δ*genP* accumulated mainly **1**, **2** and C6’-*epi*-**2**, while the double mutant Δ*genP*Δ*genK*, in which the 6’-methylated branch has been blocked due to the absence of *genK*, only produced **1** (Figs. 2a, b, Supplementary Figs. 1a, b, 2a). Δ*genB3* accumulated **1** and **2** (as previously reported^15^) as well as phosphorylated **1** and **2**, JI-20A-P (**3**) and JI-20Ba-P (**4**) (Fig. 2c, Supplementary Fig. 1c, d). Double mutant Δ*genK*Δ*genB3* produced only the 6’-unmethylated intermediates **1** and **3** (Fig. 2d, Supplementary Fig. 2b). These results not only confirm the function of GenP but also strongly implicate the role of GenB3 in the further metabolism of **3** and **4**. After incubation with a recombinant GenB3, **3** and **4**, purified from *in vitro* phosphorylation of **1** and **2** using GenP and ATP, were converted into 6’-deamino-6’-oxo-sisomicin (oxo-sisomicin, **5**) (Supplementary Fig. 1e) and 6’-deamino-6’-oxo-verdamicin (oxo-verdamicin, **6**), respectively, based on LC-ESI-HRMS and NMR analyses (Figs. 3a, b; Supplementary Figs. 1f, 3 and Supplementary Table 1), providing direct evidence that GenB3 catalyzes the removal of the 3’-phosphate and 4’-hydroxyl groups of **3** and **4** (Fig. 1). During the preparation of this manuscript, Zhou et al.^25^ published a paper on GenB3 with similar results to those described in this paper.

**Fig. 2.**
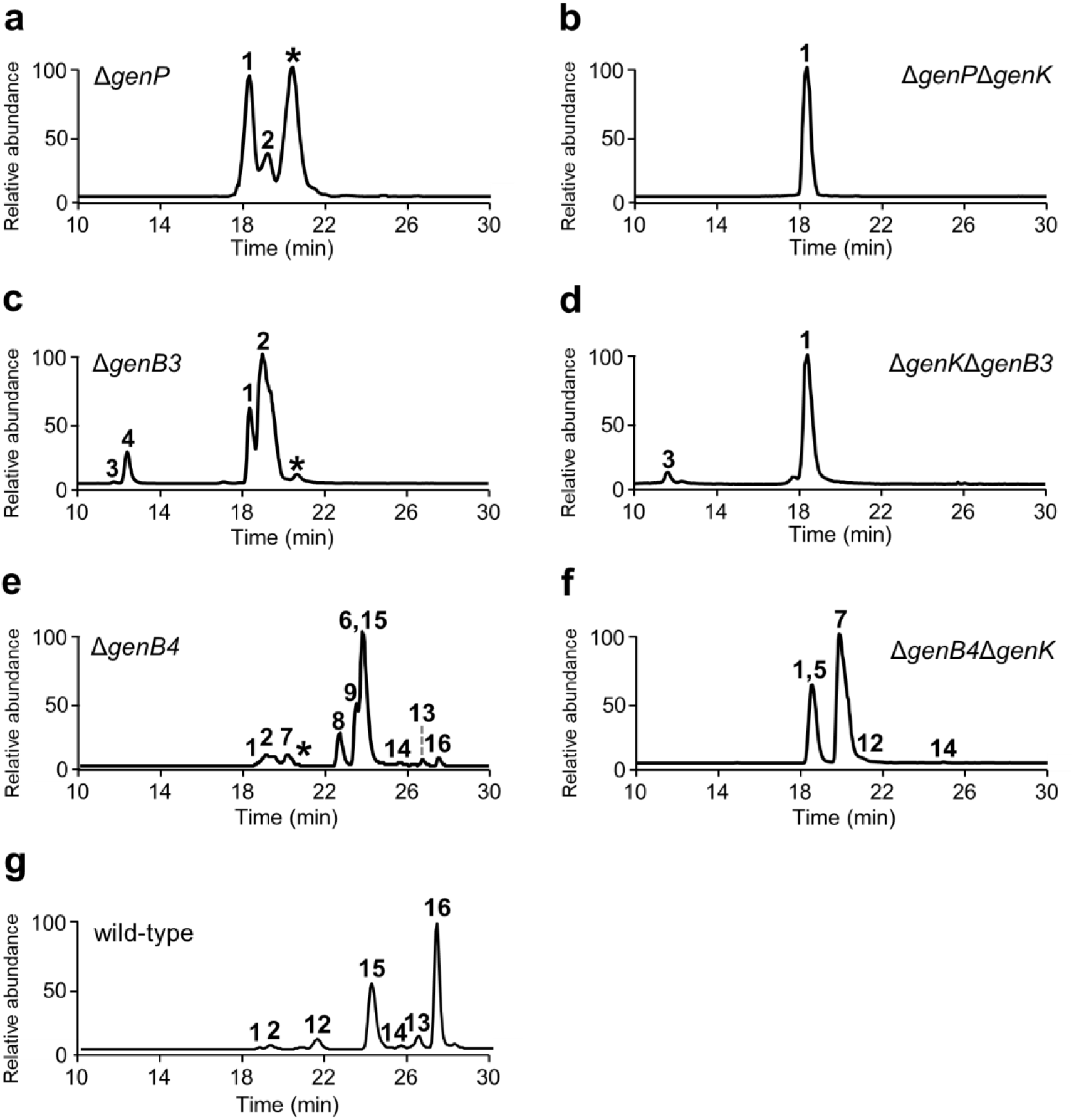
LC-ESI-HRMS analysis of gentamicin-related compounds produced *in vivo* by mutant strains. Extracted ion chromatogram traces of products from **a**, Δ*genP*; **b**, Δ*genP*Δ*genK*; **c**, Δ*genB3*; **d**, Δ*genK*Δ*genB3*; **e**, Δ*genB4*; **f**, Δ*genB4*Δ*genK* and **g**, wildtype. The star indicates JI-20Bb, which is a C6’-epimer of JI-20Ba (**2**).

**Fig. 3.**
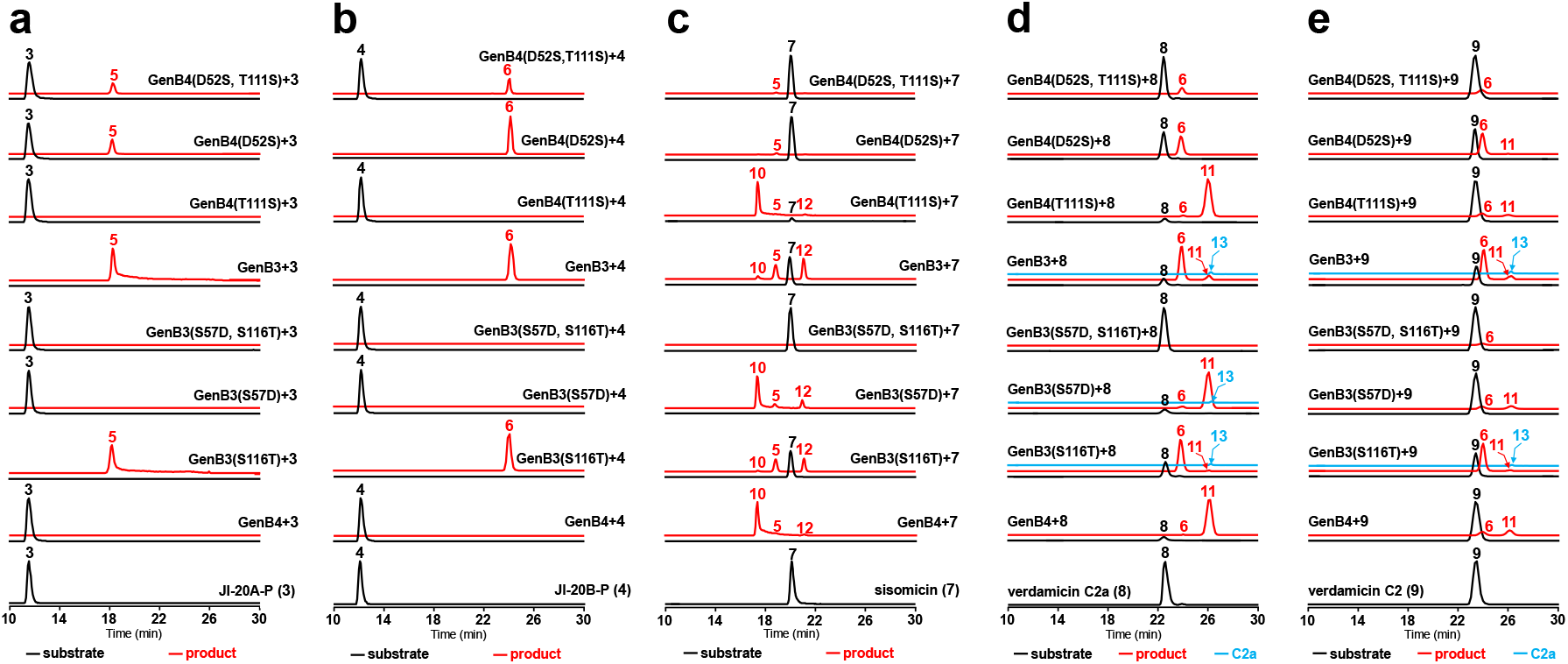
LC-ESI-HRMS analysis and comparison of the catalytic activity of GenB3, GenB4 and their mutants *in vitro*. Substrates used were: **a**, JI-20A-P (**3**); **b**, JI-20Ba-P (**4**); **c**, sisomicin (**7**); **d**, verdamicin C2a (**8**); and **e**, verdamicin C2 (**9**). Black and red lines are extracted ion chromatograms of the substrates and products, respectively. C2a (**13**) is particularly indicated by blue lines as its retention time overlapped with oxo-C2a (**11**).

We also integrated *genB3* and *genP*, coupled with *gmrA* for resistance, into the chromosome of strain ΔBN^17^, in which most of the gentamicin biosynthetic gene cluster had been deleted, for feeding experiments (Supplementary Fig. 4a). When **2** was fed to ΔBN::*genB3-genP-gmrA*, **6** was detected (Supplementary Fig. 5a). In contrast, when **1** was fed to ΔBN::*genB3-genP-gmrA*, no **5** but a trace amount of its transamination product sisomicin (7) was detected (Supplementary Figs. 1g, 5b). These experiments demonstrated that GenP phosphorylates **1** and **2** to generate **3** and **4** for GenB3. GenB3 catalyzes successive eliminations of the 4’-hydroxyl and 3’-phosphate groups resulting in a double bond at C-4’,5’ and an imine at C-6’, which is hydrolyzed giving rise to **5** and **6** (Fig. 1).

### The double bond at C-4’,5’ is reduced through double bond shift and hydrolysis by GenB4 followed by transamination by GenB3 or GenB2

GenB4 has been shown to be involved in the last step of gentamicin C complex biosynthesis and is involved in reduction of the C-4’,5’ double bond^15,18^. In the culture extract of double mutant Δ*genB4*Δ*genK* (Supplementary Fig. 2c), **5** was readily detectable and **7** was present in a much higher abundance (Fig. 2f). In contrast, only trace amounts of C1a (**12**) and C2b (**14**) (Supplementary Figs. 1h, i) were observed. Similarly, Δ*genB4* produced significant amount of **6**, verdamicin C2a (**8**) and verdamicin C2 (**9**) (Fig. 2e, Supplementary Figs. 6, 1j, k and Table 2). The chirality at C-6’ of **8** and **9** was verified through LC-MS comparison with synthetic standards kindly supplied by Prof. S. Hanessian (Supplementary Fig. 7).^26^ The ratio of gentamicin C components over other intermediates was much lower in Δ*genB4* compared to the wild-type strain. The significant decrease of C complex components in the absence of GenB4 is consistent with the notion that GenB4 is largely responsible for the C-4’,5’ double bond reduction. The presence of small quantities of Ccomplex components produced in Δ*genB4*, however, implies that the activity of GenB4 can be partially substituted by other enzyme(s).

To further understand the activity of GenB4, *in vitro* assays of purified recombinant GenB4 were performed using **6**, **7**, **8** and **9** as substrates. GenB4 robustly consumed **7** and produced a new compound *(m/z* 449.2600 and 467.2699), which was confirmed by MS/MS fragmentation to be 6’-deamino-6’-oxo-C1a (oxo-C1a, **10**) and its hydrate (6’,6’-diol), and small amounts of **12** and **5** (Fig. 3c, Supplementary Fig. 1l). GenB4 also efficiently converted **8** to 6’-deamino-6’-oxo-C2a (oxo-C2a, **11**) (*m/z* 463.2747) (Fig. 3d, Supplementary Fig. 1m). Thus, GenB4 catalyses 4’,5’-reduction and concomitant 6’-oxidation/deamination on its substrates. It is noteworthy that **6** was not utilized by GenB4 (Fig. 4a) and that **9**, the epimer of **8**, was only converted at low efficiency (Fig. 3e), strongly suggesting that the activity of GenB4 requires the presence of the C-6’ amino group and is influenced greatly by the stereochemistry at C-6’.

**Fig. 4.**
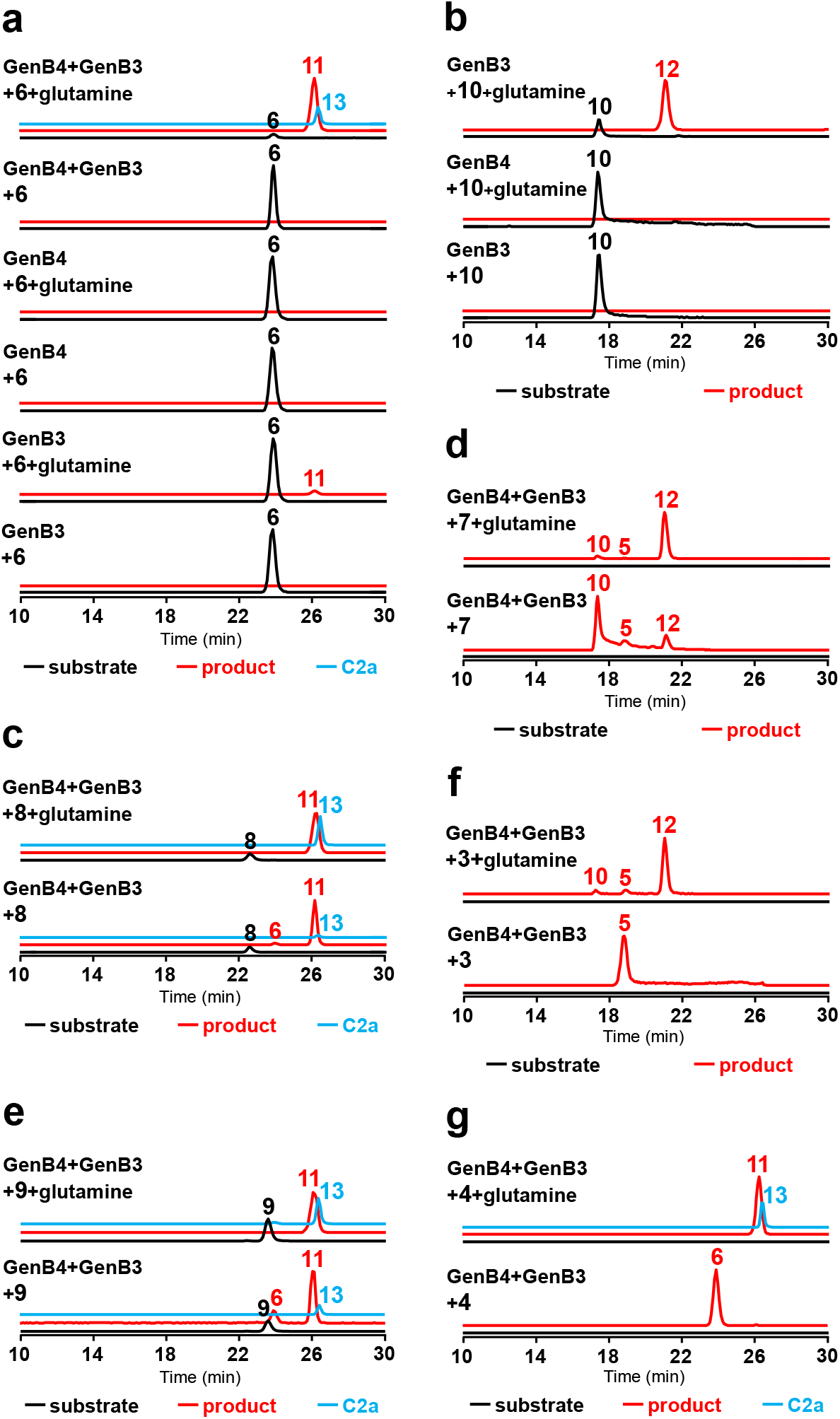
LC-ESI-HRMS analysis of the influence of amino donor L-glutamine on the generation of C1a (12) and C2a (13) *in vitro*. Substrates used were: **a**, oxo-verdamicin (**6**); **b**, oxo-C1a (**10**); **c**, verdamicin C2a (**8**); **d**, sisomicin (**7**); **e**, verdamicin C2 (**9**); **f**, JI-20A-P (**3**); and **g**, JI-20Ba-P (**4**). Black and red lines are extracted ion chromatograms of the substrates and products, respectively. C2a (**13**) is particularly indicated by blue lines as its retention time overlapped with oxo-C2a (**11**).

Furthermore, feeding experiments were performed to investigate whether GenB4 can reduce sisomicin and verdamicin to produce gentamicin C components *in vivo*. When 6’-unmethylated sisomicin **7** was fed to ΔBN::*genB4-gmrA* (Supplementary Fig. 4b), **10** was the most abundant species accumulated, although a trace amount of **5** and a small amount of **14** w4ere also detected (Supplementary Fig. 5c). Feeding the 6’-methylated verdamicin **8** to *ΔBN::genB4-gmrA* yielded a very low level of C2a (**13**) (Supplementary Figs. 1n, 5d) with **11** being the major product. As in our *in vitro* study, the conversion was not significant when **9** was fed (Supplementary Fig. 5e).

GenB3 was also tested as a candidate enzyme responsible for the small amount of gentamicin C components produced in Δ*genB4*. Incubation of GenB3 with **7** produced a mixture containing **5** and **12** at a similar level with a relatively low yield of **10** (Fig. 3c), indicating that the reaction might have followed a disproportionation mechanism, in which the amino group from some molecules of **7** is used to transaminate **10** produced from other molecules of **7**, giving **12**. In contrast, the main product of GenB3-catalyzed transformation of **8** and **9** was **6**, which was also accumulated in Δ*genB4*, and the yield of **11** and **13** are both low (Figs. 3d, e). These results indicate that the efficiency of GenB3-catalyzed 4’,5’-double bond reduction in the 6’-methylated branch of the pathway is lower than that in the 6’-unmethylated branch *in vitro*, showing the influence of the C6’ methyl group.

### GenB3 is responsible for C-6’ transamination following both didehydroxylation and double bond reduction

Examining the above-mentioned results on activities of GenB3 and GenB4 revealed a missing link: transfer of an amino group to C-6’ of the products of GenB3 (**5** and **6**) to generate sisomicin **7** and verdamicins (**8** and **9**), the substrates for GenB4. GenB3 was tested as a candidate enzyme for this amination since it has been shown that GenB3 is able to catalyze C-6’ amination on earlier intermediates 6-DOG and 6-DOX using L-glutamine as an amino donor^15^. Unexpectedly, in the presence of L-glutamine, GenB3 only catalyzed production of a small amount of **11** from **6**, but no **8** or **9** were detectable (Fig. 4a). This result suggests that the reversible transamination is in favour of a ketone group at C-6’ and GenB3 may be inhibited by low levels of its C-6’ aminated product, similarly to the transamination performed by GenS2^16^. In a one-pot reaction containing GenB3, GenB4, **6** and L-glutamine, transformation of **6** to both **11** and **13** were observed (Fig. 4a), suggesting that alleviation of the product inhibition on GenB3 by GenB4-catalyzed 4’,5’-reduction of verdamicins allowed the biosynthetic process to proceed forward.

Feeding **6** to ΔBN::*genB3-gmrA* only yielded a small amount of **8** (Supplementary Fig. 5f), while feeding **8** resulted in production of **6** in a significant amount, again demonstrating that GenB3 catalyzed-transamination favours deamination over amination. As expected, **6** was converted to **13** when fed to ΔBN:: *genB4-genB3-gmrA* (Supplementary Figs. 4c, 5f).

Oxo-C1a **10** and oxo-C2a **11** were the dominant products of GenB4-catalyzed transformation of sisomicin and verdamicins (Figs. 3c-e), indicating another enzyme catalysing C-6’ amination is needed in order to convert **10** and **11** into **12** and **13**, respectively. GenB3 was tested as a candidate for this amination since Δ*genB1*Δ*genB2* harbouring *genB3* produced a small amount of gentamicin C components^15^. Results shown in Fig. 4 demonstrate that efficient conversion of **10** to **12** by GenB3 (Fig. 4b) and of **7**, **8** and **9** to **12** or **13** by combined activities of GenB3 and GenB4 were efficiently achieved only in the presence of L-glutamine (Figs. 4c-e), confirming the role played by GenB3 and an amino donor in this last amination step of gentamicin biosynthesis.

Our *in vivo* feeding studies showed that GenB2 also displayed C-6’ amination activities on the intermediates produced. Both ΔBN::*genB3-gmrA* and ΔBN::*genB2-gmrA* converted **10** to **12** and **14** (Supplementary Figs. 4d, e, 5g). Feeding **7** to ΔBN::*genB4*-*genB3-gmrA* and ΔBN::*genB2-gmrA-genB4* (Supplementary Figs. 4c, f, 5c) resulted in production of **12** and **14** by both strains. Conversion of both **8** and **9** to **13** was observed with both ΔBN::*genB4-genB3-gmrA* and ΔBN::*genB2-gmrA-genB4* (Supplementary Fig. 5d), although the latter strain also produced C2 (**15**) and C1 (**16**) (Supplementary Figs. 1o, p) and the conversion of **9** was less efficient. (Supplementary Fig. 5e). In all cases, GenB3 appeared to have higher activity than GenB2 in the transamination of **10** and **11**.

Finally, as further confirmation for the roles played by GenP, GenB3 and GenB4 in the biosynthetic pathway from **1** and **2** to gentamicin C components, reconstitution of the dideoxygenation process *in vitro* was carried out: GenB3 and GenB4 were assayed starting from **3** and **4;** and **1** and **2** were fed to ΔBN::*genB3-genP-genB4-gmrA* (Supplementary Fig. 4g). In the absence of L-glutamine, the coupled activities of purified GenB3 and GenB4 transform **3** and **4** to **5** and **6**, respectively (Figs. 4f, g). When an excess of L-glutamine was added to the reaction **12** and **13** became the main products. Moreover, **1** and **2** fed to ΔBN::*genB3-genP-genB4-gmrA* were converted to **12** and **13**, respectively (Supplementary Figs. 4g, 5a, b). JI-20Ba (**2**) and its 6’-epimer JI-20Bb (Supplementary Fig. 1q) were proposed to be the precursors of C2a and C2^19^. In our *in vitro* assays, both of them were phosphorylated by GenP (Supplementary Figs. 8, 1r). However, only JI-20Ba-P (**4**) was taken by GenB3 as a substrate.

### Crystal structures of GenB3 and GenB4

The observation that GenB3 and GenB4, despite sharing very high sequence homology, catalyse different reactions prompted us to obtain crystal structures of the two proteins in order to understand the structural basis for their catalytic activities. Crystal structures of GenB3 and GenB4 with their prosthetic group PLP (GenB3-PLP, GenB4-PLP) and GenB4-PLP co-crystallised with sisomicin **7** were solved with nearly atomic resolution (Supplementary Table 3).

GenB3 and GenB4 structures are highly similar with an RMSD of 0.47 Å (Supplementary Figs. 9a, 10a), but are divergent from GenB1 (RMSD of 2.05 Å) sharing only about 27% identity (Supplementary Figs. 9b, 10b). GenB3 and GenB4 both crystallized as dimers that have the characteristic type I transaminase fold composed of three domains: an N-terminal domain, a C-terminal domain and a PLP binding domain (Figs. 5a, b). The two protomers form a dimer mainly through hydrophobic interactions. Superposition of the two protomers in the asymmetric unit shows high similarity (RMSD of 0.21 Å for GenB3 and 0.34 Å for GenB4, respectively). The PLP and the active site are located at the interface of the protomers. In GenB4 and GenB3, PLP forms a Schiff base with Lys238 and Lys243, respectively (Supplementary Figs 11a, b). In GenB4, the pyridine ring nitrogen of PLP is hydrogen bonding with the side chain of Asp210 and the main chain carbonyl of Tyr135. A hydrogen bond network connects the phosphate of PLP with the side chains of Thr111 and Thr267 from the other protomer as well as with the main chain nitrogen atoms of Gly110 and Thr111. Additionally, Trp138 and three residues of the neighbouring protomer, His107, Tyr268 and Thr267, also form hydrogen bonds with the PLP-phosphate via two water molecules (Fig. 5c). These interactions between PLP and GenB4 were conserved in GenB3 except that the Thr111 is replaced by a serine.

**Fig. 5.**
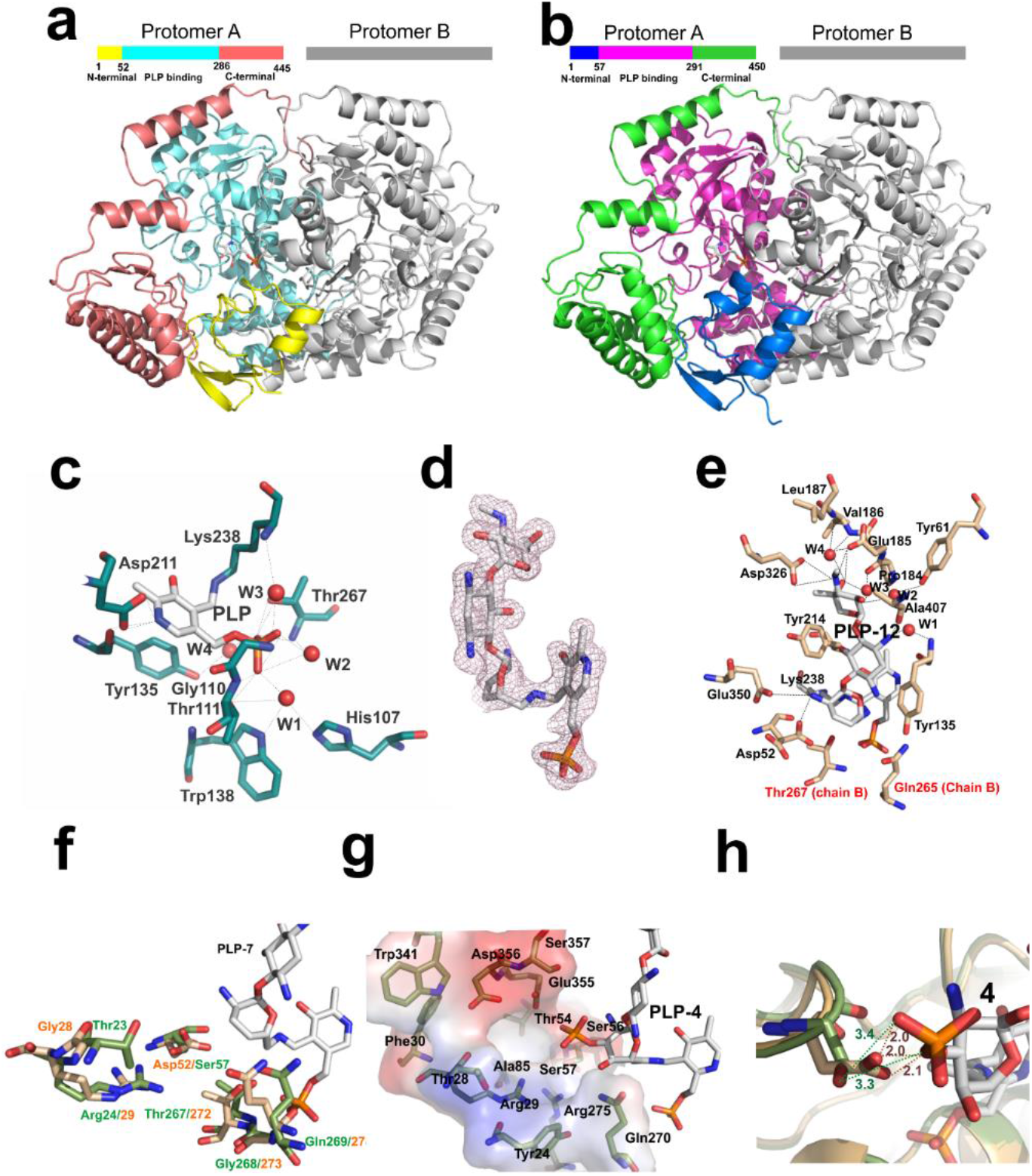
3D structures of GenB3 and GenB4. **a, b**, The overall structures of GenB4 and GenB3, respectively. The structure of the dimer is represented. The colours in protomer A indicate the different domains, as shown with the bars on the top of each structure. The second protomer of both GenB4 and GenB3 is represented in grey. **c**, PLP binding site of GenB4-PLP. The lines represent the hydrogen bond interactions between the protein amino acid residues (carbons in green) and the PLP (carbons in white). The red spheres are water molecules. **d**, Electron density contours for the external aldimine of PLP-**12** in GenB4-PLP-**12**. **e**, Residues at the active site of GenB4-PLP-**12** (carbons in beige) and the external aldimine of PLP-**12** (carbons in white). Dotted lines indicate hydrogen bond interactions and water molecules are shown as red spheres. **f**, Major differences of residues in the active site cavity of GenB3 and GenB4. The amino acid residues from GenB3 are represented in green and from GenB4 are in beige. PLP-**12** is from GenB4-PLP-**12** and is shown with carbons in white. **g**, The hypothetical binding site for the phosphate group of **4** in GenB3. The amino acid residues are shown as sticks. The contours are the electrostatic potential surface for GenB3. Red and blue colors indicate negatively charged and positively charged regions, respectively. **h**, Representation of hypothetical binding of **4** in GenB3 and the superposition with GenB4. The Asp52 (carbons in beige) and Ser57 (carbons in green) from GenB4 and GenB3, respectively are shown in sticks. The distances represented in orange are from GenB4 Asp52 side chain to the phosphate group and the distances in green are from GenB3 Ser57 side chain to the phosphate group of **4.**

The crystals obtained from GenB4-PLP-**7** show electron density for an aminoglycoside in both active sites and binding of this aminoglycoside does not induce significant changes in the overall structure of GenB4 (RMSD of 0.22 Å) (Supplementary Fig. 12), although conformational changes of a few residues in the active sites are seen. However, modelling of possible structures into the electron density revealed that the aminoglycoside no longer has the 4’,5’-double bond of **7**. The structure that fits best is gentamicin C1a **12**. This would be formed by the expected GenB4-catalysed conversion of **7** into **10** followed by transamination of **10** to **12** by GenB4 with residual **7** as the amino donor. In one of the protomers of the GenB4-PLP-**12** dimer, a *gem*-diamine formed by PLP with Lys238 and **12** was observed, which is an intermediate between the internal aldimine with Lys238 and the external aldimine with **12** (Supplementary Figure 13). In the second protomer, Lys238 has moved away from PLP to form a hydrogen bond with Thr267 and the PLP forms the external aldimine with **12** (Fig. 5d, Supplementary Fig. 14). Therefore the two active sites of the dimer are at different catalytic stages. Interestingly, in the protomer where the *gem*-diamine is found, the side chain of Asp52 displays double conformations with one hydrogen bonding to Arg270 and Glu350 as observed in GenB4-PLP, and the other flipping about 90° to form a hydrogen bond with the 2’-NH_2_ of **12**, suggesting that Asp52 and associated water molecules may play an important role in substrate binding and/or enzymatic activity (Supplementary Fig. 14b). Other interactions between **12** and GenB4 include hydrogen bonds between the 3”-methylamine group of ring III and the side chains of Glu185 and Asp326, a weak π-stacking between the 2-DOS ring and Tyr135, and a hydrogen bond of the 1-NH_2_ with Ala407 main chain carbonyl. Several water molecules in the active site groove form bridging hydrogen bonding interactions between **12** and GenB4 residues (Figs. 5e).

The electrostatic surface potential of GenB4 shows a negatively charged cavity favorable for accommodating cationic aminoglycoside substrates, while the cavity in GenB3 appears to be slightly more positive (Supplementary Fig. 15). Superposing GenB4-PLP-**12** with GenB3-PLP structures reveals a few residues around the substrate-binding pocket that are different between the two proteins (Fig. 5f), In order to understand if these differences are related to substrate specificities of the two enzymes, we have modelled **4** into GenB3-PLP. The phosphate group is accommodated in a pocket formed by Tyr24, Arg29, Ser57, Ser357, Glu355 and Asp356 together with Gln270 and Arg275 from another protomer (Fig. 5g). In particular, the phosphate group forms hydrogen bonds with side chains of Ser57 and Glu355 and those of Arg29, Ser57 and Arg275 through water molecules, which may serve to stabilize the GenB3-PLP-**4** intermediate as well as to make the 3’-phosphate a better leaving group (Fig 5g). The side chain of Gln270 hydrogen bonds to the 4’-hydroxyl, possibly playing a part in the 4’,5’-dehydration step. All the residues in this region are conserved in GenB4 except that Ser57 is replaced by Asp52. An attempt to model **4** into GenB4 puts the 4’-phosphate group in a position too close (2 Å ~ 2.1 Å) to the side chain of Asp52, where repulsion between the negatively charged carboxyl side chain of Asp52 and the phosphate moiety should prevent binding of **3** and **4** in the pocket (Fig. 5h), which could explain why GenB4 does not accept them as substrates.

### Ser57 of GenB3 and Asp52 of GenB4 are crucial for their catalytic activities

Analysis of the crystal structures of GenB3 and GenB4 has allowed us to identify two residues in the active sites that are different between these two otherwise highly homologous enzymes: Ser57 and Ser116 in GenB3 and their equivalents Asp52 and Thr111 in GenB4. To investigate whether these differences provide structural bases for their different activities, residue-swapping between GenB3 and GenB4 was carried out by site-directed mutagenesis creating four single mutants GenB3(S57D), GenB3(S116T), GenB4(D52S) and GenB4(T111S) and two double mutants GenB3(S57D/S116T) and GenB4(D52S/T111S). Purified mutant enzymes were characterized by *in vitro* enzymatic assays using the wild-type enzymes as controls.

As shown in Figs. 3a and 3b, GenB4(D52S) has acquired activity to catalyze didehydroxylation on **3** and **4**, but the activity possessed by wild-type GenB4 to reduce the C-4’,5’ double bond was impaired. When **7**, **8** and **9** were used as substrates, GenB4(D52S) behaved more like wild-type GenB3 producing mainly C-6’-deaminated intermediates (Figs. 3c-e). On the other hand, S57D mutation in GenB3 resulted in complete loss of the activity for catalyzing didehydroxylation on both **3** and **4** (Figs. 3a, b). GenB3(S57D), however, displayed significant activity in reducing the C-4’,5’ double bond of **7** and **8**, although **9** appeared to be a poor substrate for this activity (Figs. 3c-e). GenB3(S116T) and GenB4(T111S) showed similar activities to their wild-type enzymes (Fig. 3). As expected, GenB3(S57D/S116T) and GenB4 (D52S/T111S) showed similar activities to GenB3(S57D) and GenB4 (D52S), respectively (Fig. 3). These results demonstrate that the key residue dictating the enzymic activity of GenB3 is Ser57 and of GenB4 is Asp52.

## Conclusion

In answer to the question of how the 3’- and 4’-hydroxyls of **1** and **2** are removed to produce gentamicin C complex during the later stages of gentamicin biosynthesis, experimental evidence obtained in this study demonstrates that GenP, GenB3 and GenB4 together are responsible for catalyzing the didehydroxylation process, in which GenB3 and GenB4 exhibit types of activity that have not been reported so far for known PLP-dependent enzymes, in addition to the transaminase activity predicted by sequence homology. Catalytic mechanisms for GenB3 and GenB4 are proposed in Fig. 6. C-3’-phosphorylation of **1** and **2** by GenP generates substrates for GenB3. GenB3 catalyzes elimination of the 4’-hydroxyl and then of the 3’-phosphate, which results initially in 3’,4’ and 5’,6’ double bonds. Imine exchange with the active site lysine liberates an enamine, which protonates on C-3’, giving an imine with a 4’,5’-double-bond. Hydrolysis of the imine gives **5** and **6**. The mechanism is similar to that of methionine γ-lyase, also a PLP-dependent enzyme^27^, but incorporates the additional elimination of a second leaving group (the 3’-phosphate). We know of no other PLP-dependent enzyme that catalyzes the elimination of a leaving group at this distance from the reactive amino group. The C6’ carbonyls of **5** and **6** are converted to amine form by GenB3 with glutamine as an amino donor, yielding **7** and **8**. The 4’,5’-reductions with concomitant oxidations at C-6’ are performed on **7** and **8** by GenB4, or less efficiently by GenB3, by protonation at C-4’ of the quinonoid intermediate, thus shifting the 4’,5’ double bond to the 5’,6’ position. Imine exchange again liberates an enamine, which protonates at C-5’, giving an imine.

**Fig. 6.**
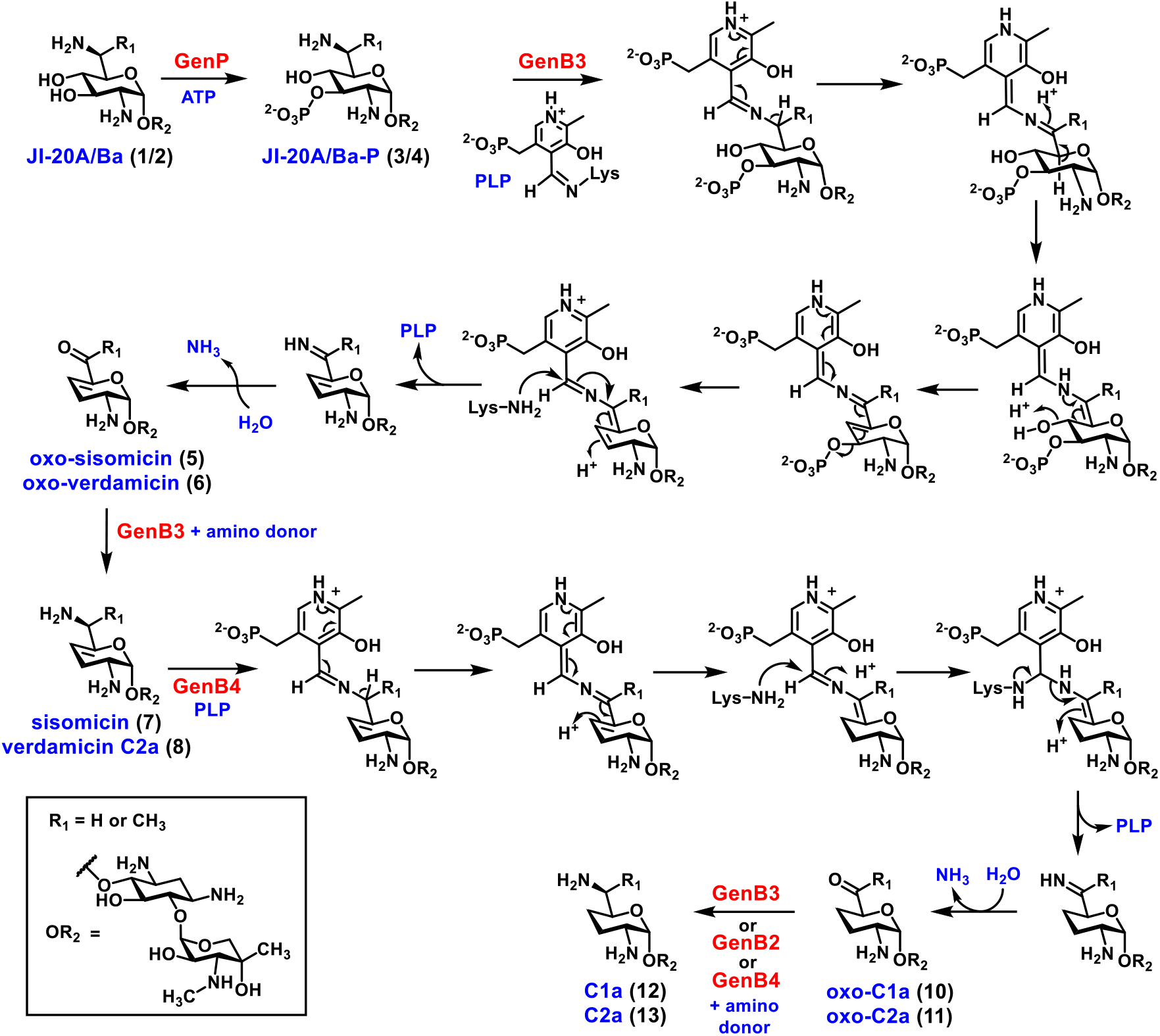
Proposed mechanisms for the conversion of JI-20A/Ba (1 and 2) to gentamicin C1a (12) and C2a (13) catalyzed by coupled activities of GenP, GenB3 and GenB4.

Hydrolysis of the imine generates **10** and **11**. Yet another transamination of the C-6’ carbonyls of **10** and **11** by GenB3 (or GenB2) with an amino donor leads to gentamicins C1a (**12**) and C2a (**13**), respectively. In *in vitro* experiments using sisomicin **7** or verdamicin C2 **8** as substrates, these compounds can act as the amino donor if no other suitable amino donor is included.

Analysis of the crystal structures reveals that only two of the residues lining the active sites differ between GenB3 and GenB4. Of particular interest are Ser57 in GenB3 and Asp52 in GenB4. A hydrogen bond is predicted between Ser57 and the 3’-phosphate of **4**, which could be crucial for stabilizing the enzyme-substrate intermediate and later helping the departure of the phosphate group. We speculate that replacing the Ser57 with an Asp would interfere with the binding of **3** and **4** due to electronic repulsion. The binding of **12** to GenB4 induced a significant flipping of the side chain of Asp52 to form a hydrogen bond with the 2’-NH_2_ of **12**, which possibly plays a role in the double bond migration during the catalysis. In support of the hypotheses, swapping the residues between the two enzymes has allowed the corresponding switching of functions, with a loss of or a decrease in the original activities.

## Methods

Methods describing gene cloning, gene deletion and complementation, fermentation of strains and feeding experiments, isolation and characterization of compounds, LC-ESI-HRMS analysis, protein overexpression and purification, enzymatic assay, crystallization and related analysis are described in full details in Supplementary Information.

## Data availability

Coordinates and structure factors have been deposited with the Protein Data Bank with accession codes 7LLD and 7LLE and 7LM0. All other data are available from the authors on reasonable request.

## Acknowledgments

This work was supported by the National Key R&D Program of China (2018YFA0903200) and the Funds for International Cooperation and Exchange of the National Natural Science Foundation of China (31920103001) to Y.S., the São Paulo Research Foundation (FAPESP) (2011/15971-3, 2014/50324-0, 2015/09188-8 and 2018/00351-1) to M.V.B.D., the fellowship from FAPESP (14/07843-6) to P.S.B., the Medical Research Council (MRC) Grants G1001687 and MR/M019020/1 to P.F.L. and the MRC postgraduate studentship (1343325) to A.R. We thank Professor Stephen Hanessian (Université de Montreal, Canada) for kindly providing synthetic verdamicin C2 and verdamicin C2a.

## Author contributions

Y.S., P.F.L. and M.V.B.D. designed the study. S.L., J.G., C.H. and X.J. performed gene deletion, complementation and feeding experiments. S.L., F.H., M.V.B.D., J.G. and A.R. performed protein expression, purification, compound isolation and *in vitro* experiments. G.S., S.L. and C.H. performed compound characterization. P.S.B and M.V.B.D. performed protein crystallization and solved the crystal structures. All authors analyzed and discussed the results. S.L., F.H., M.V.B.D., Y.S., F.J.L. and P.F.L. prepared the manuscript.

## Additional information

Supplementary information is available for this paper at https://doi.org/

