## Supporting information for "Mechanistic insights into dideoxygenation in gentamicin biosynthesis"

### Supplementary Information

**Bacterial strains, chemicals, and culture Conditions.** *E. coli* DH10B was used as a cloning host and *E. coli* ET12567/pUZ8002 for intergeneric conjugation between *E. coli* and *Micromonospora*[1]. *Micromonospora echinospora* ATCC15835 (wild-type), Δ*genP* and Δ*genK*[2,3] were used for creating single or double in-frame deletion mutants and as the source of *gen* genes. Restriction endonucleases, Phusion High-Fidelity Master Mix with GC-buffer, and T4 DNA ligase were purchased from New England Biolabs. Oligonucleotide primers were synthesized by GenScript and Tsingke. DNA sequencing of PCR products was performed by GenScript or by the Department of Biochemistry DNA Sequencing Facility, University of Cambridge. DIG DNA labeling and detection kits were purchased from Roche. For standards, sisomicin (**7**) and gentamicin C complex were purchased from Sigma-Aldrich.

*M. echinospora* strains were grown in ATCC172 medium (soluble starch 2%, glucose 1%, yeast extract 0.5%, N-Z amine type A 0.5%, $CaCO_3$ 0.1%) for chromosomal DNA isolation and preparation of mycelium. *E. coli* strains were maintained in 2×TY (tryptone 1.6%, yeast extract 1.0%, NaCl 0.5%) media at 37°C with the appropriate antibiotic selection at a final concentration of 100 μg/mL for ampicillin, 25 μg/mL for kanamycin, and 25 μg/mL for chloramphenicol.

**Fermentation and feeding experiments**. *M. echinospora* ATCC15835 and its mutants were cultured in two stages. Seed culture was maintained in liquid ATCC172 medium at 28°C with shaking at 220 rpm for 2 days before being inoculated into 30 mL of F50 fermentation medium[4] with 5% inoculum, then incubated at 28°C with shaking at 220 rpm for 5 days. For feeding experiments in vivo, filter-sterilized compounds were added to the medium at 20 μg/mL of final concentration when the seed culture was inoculated into the fermentation medium. For preparing intermediates on large scale, the second stage was performed in 20 L of F50 fermentation medium[4] shaken at 220 rpm and at 28°C for 5 days.

**Construction of gene disruption plasmids.** For in-frame deletion of a target gene by homologous recombination, two DNA fragments flanking the targeted sequence were amplified from the genomic DNA of *M. echinospora* ATCC15835 by using two pairs of primers (Supplementary Table 4). The PCR products were each cloned into pUC18, then inserted together into the *Streptomyces*-*E. coli* shuttle vector pYH7[5] to obtain the gene disruption plasmids (Supplementary Table 5), which was verified by sequencing.

**Targeted in-frame gene deletion.** To create in-frame deletion mutants, the corresponding plasmids constructed for gene knock-out were introduced into the appropriate hosts by conjugation and mutants screened by the same method as described before[4]. The desired in-frame deletion mutants were identified by PCR (Supplementary Tables 4, 6).

**Gene complementation.** For gene complementation, target genes or sequences were cloned from *M. echinospora* ATCC15835 genomic DNA, then inserted into vector pWHU77[2] between *Nde*I and *Eco*RI sites. When complementing multiple discrete genes, all fragments and the linearized vector were integrated into one pot by Gibson assembly kit (New England Biolabs) to generate the recombinant plasmids (Supplementary Table 5). After sequence confirmation, these plasmids were introduced individually into ΔBN by conjugation as described previously[2]. Complemented exconjugants were verified on A medium (soluble starch 1%, corn steep powder 0.25%, yeast extract 0.3%, $CaCO_3$ 0.3%, $FeSO_4$ 0.0012%, agar 3%) containing thiostrepton (25 μg/mL) and confirmed by PCR (Supplementary Table 4).

**Site-directed mutations of GenB3 and GenB4.** Site-directed mutations of GenB3 and GenB4 were generated using the QuikChange (Stratagene) method (Supplementary Table 7). The *genB3* and *genB4* genes, respectively, were inserted into a pUC18 vector between the *Nde*I and *Bam*HI sites as the template. PCR amplifications were carried out using HF Phusion DNA polymerase (NEB) with 30 cycles of denaturation at 98°C for 10 sec, annealing at 60°C for 30 sec, and extension at 72°C for 2 min followed by a final extension at 72°C for 10 min. The resulting PCR products were digested with *Dpn*I at 37°C for 1 h to remove the template before being introduced into *E. coli* NovaBlue cells (Novagen). Plasmids bearing the desired mutation identified by DNA sequencing were isolated from the transformants, digested with restriction enzymes *Nde*I and *Bam*HI, purified by gel extraction, and inserted into plasmid pET28a(+). The inserts of the recombinant plasmids were verified by DNA sequencing.

**Extraction of aminoglycosides.** Cultures of mutants of *M. echinospora* ATCC15835 or cell-free product were adjusted to pH 2.0 with $H_2SO_4$ and agitated for 2 h. The supernatant after centrifugation was filtered through Whatman filter paper and agitated for 2 h with DOWEX 50 WX8-200 ion-exchange resin (1 g for 30 mL broth or 3ml cell-free product) that was preconditioned with acetonitrile followed by Milli-Q water. The resin was put in a column then washed with Milli-Q water (6 column volumes) and eluted with 1 M $NH_4OH$ (6 column volumes). The eluate was mixed with #711 anion ion-exchange resin (1 g for 30 mL broth or 5 mL cell-free product) and agitated for 1 h, then the supernatant was freeze-dried and redissolved in Milli-Q water (0.3 ml concentrated solution was equivalent to 30 mL broth or 3 mL cell-free product), and filtered through 0.22 μm microporous membrane.

**LC-ESI-HRMS analyses.** LC-ESI-HRMS analysis was performed on Thermo Electron LTQ-Orbitrap XL fitted with Phenomenex Luna C18 column (250×4.6 mm) at flow rate 0.4 mL/min, using a mobile phase of (A) 0.2% trifluoroacetic acid (TFA) in $H_2O$ (adjusted to pH 2.0 with $NH_4OH$) and (B) 100% $CH_3CN$; the gradient for separation of gentamicin complex and intermediates was: 0-14 min 2% B to 6% B, 14-16 min 6% B to 8% B, 16-20 min 8% B to 15% B, 20-25 min 15% B, 25-27 min 15% B to 90% B, 27-31 min 90% B (in this section, the flow rate was set 0.6 mL/min), 31-31.5 min 90% B to 2% B, 31.5-35.5 min 2% B. MS/MS analyses were carried out in positive ionization mode with 35% relative collision energy.

***In vitro* reactions with GenP, GenB3, GenB4 and their mutants.** Every involved gene or its mutant was inserted in pET28a(+) and expressed with His-tag as described above. The harvested cells were opened through a high-pressure homogenizer in a buffer containing 25 mM Tris-HCl, 300 mM NaCl and 25 mM imidazole (pH 7.4). After centrifugation, the reconstituted protein in the supernatant was purified using $Ni^{2+}$ ion-charged His-Bind metal chelating resin and eluted by a buffer containing 25 mM Tris-HCl, 300 mM NaCl and 300 mM imidazole (pH 7.4). The fraction with the highest protein concentration was further purified through Superdex™ 200 Increase 10/300 GL in ÅKTA Purifier (GE Healthcare) using a mobile phase containing 25 mM Tris-HCl, 300 mM NaCl and 10% glycerin (pH 7.4). The purified proteins were concentrated and stored in the same buffer.

For in vitro assays, the reactions were performed in 150 μL buffer containing 50 mM Tris-HCl (pH 7.4). The concentration of each component in most reactions is: aminoglycoside about 0.02 mg/mL; GenB3, GenB4 and their mutants 1 mg/mL; glutamine 1 mg/mL. Aminoglycosides were adjusted to about pH 7.0 before adding to reactions. Reaction mixtures were incubated at 28°C for 8 h when using oxo-verdamicin (**6**), sisomicin (**7**), verdamicin C2a (**8**), verdamicin C2 (**9**) or oxo-C1a (**10**) as substrates, or overnight when using JI-20A (**1**), JI-20Ba (**2**), JI-20A-P (**3**) or JI-20Ba-P (**4**) as substrates. All reactions were quenched by the addition of 150 μL chloroform followed by vigorous vortexing. Each sample was filtered through 0.22 μm microporous membrane before detection. The JI-20Ba (**2**) and its 6'-epimer JI-20Bb were purified from 1.5 mL reaction mixture with GenB2 and JI-20Ba (**2**). For reactions involving GenP, 2 mg/mL ATP and 1 mM $Mg^{2+}$ were added. For the preparation of JI-20A/B-P (**3**/**4**), the concentration of substrate JI-20A/B (**1**/**2**) was increased to 0.2 mg/mL.

**Isolation and purification of aminoglycosides.** Isolation and purification of gentamicin related compounds from crude extracts and JI-20A/B-P (**3**/**4**) from reaction of GenP and ATP with JI-20A/B (**1**/**2**) were performed on Thermo Scientific HPLC (UltiMate 3000) fitted with an evaporative light scattering detector (ELSD, Alltech 2000ES) and a Phenomenex Synergi C18 column (250×10 mm) at a flow rate of

3.9 mL/min, using a mobile phase of (A) 0.2% TFA in $H_2O$ and (B) 100% $CH_3CN$; the gradient for separation of both gentamicin complex and related intermediates was: 0-10 min 2% B to 4.5% B, 10-10.5 min 3.5% B to 90% B, 10.5-15.5 min 90% B, 15-15.5 min 90% B to 2% B, 15.5-19 min 2% B. The temperature and gas flow of ELSD was 111°C and 2.9 L/min. JI-20Ba (**2**) and JI-20Bb were purified in mini scale from the reaction mixture of GenB2 and JI-20Ba (**2**) on Thermo Electron LTQ-Orbitrap XL fitted with Phenomenex Luna C18 column (250×4.6 mm) at flow rate 0.4 mL/min, using a mobile phase of (A) 0.2% TFA in $H_2O$ and (B) 100% $CH_3CN$; the gradient for separation was: 0-14 min 2% B to 6% B, 14-16 min 6% B to 8% B, 16-20 min 8% B to 15% B, 20-25 min 15% B, 25-27 min 15% B to 90% B, 27-31 min 90% B (in this section, the flow rate was set 0.6 mL/min), 31-31.5 min 90% B to 2% B, 31.5-35.5 min 2% B.

**NMR characterization.** The 1D ($^1$H, $^{13}$C and DEPT) and 2D (COSY, HSQC, HMBC and NOESY) NMR spectra were collected on an Agilent-NMR-VNMRS 600 spectrometer. Chemical shifts are reported in ppm, and NMR data processing was performed using MestReNova software.

**Structure Determination of oxo-verdamicin (6), verdamicin C2a (8) and verdamicin C2 (9).** The molecular formula of oxo-verdamicin (**6**) was deduced to be $C_{20}H_{36}N_4O_8$, based on the quasi-molecular ion peak at *m/z* 461.2567 $[M + H]^+$ (calc. for $C_{20}H_{37}N_4O_8^+$, 461.2606) in its ESI-HRMS spectrum (Supplementary Fig. 1f). The singlet methyl proton signals [$\delta_H$ 1.35 (s, 3H), 2.38 (s, 3H), 2.92 (s, 3H)] in the $^1$H NMR spectrum (Supplementary Fig. 3a) indicated three methyl groups. A C=C double bond and a ketone could be inferred from the $^{13}$C signals: $\delta_C$ 115.1, 148.1 and 197.3 (Supplementary Fig. 3b). Two $^{13}$C signals, $\delta_C$ 98.5, 102.6, as well as the corresponding $^1$H signals, $\delta_H$ 5.62 (s, 1H), 5.09 (d, 3.7 Hz, 1H), indicated that oxo-verdamicin (**6**) contained two glycoside residues. $^1$H-$^1$H COSY correlations of H-1/H-2/H-3/H-4/H-5 along with the HMBC correlations of H-6 with C-1 & C-5 and H-2 with C-1, C-3, C-4 & C-6 (Supplementary Figs. 3c-e) determined the fragment of 1,3-diamino-4,5,6-tri-O-cyclohexane (2-DOS). The HMBC correlations of H-1' with C-2', C-3' & C-5', H-4' with

C-2’, C-5’ & C-6’, and 6’-$CH_3$ with C-5’ & C-6’ combined with $^1$H-$^1$H COSY correlations of H-1’ with H-2’ and H-3’ with H-4’ (Supplementary Figs. 3c-e) determined the planar structure of 2’-$NH_2$-6’-oxo-$\Delta^{4',5'}$-heptose. The planar structure of another sugar residue was deduced to be 2’’,4’’-dihydroxy-4’’-methyl-3’’-methyl-amino-pentose according to HMBC correlations of H-5’’ with C-1’’ & C-4’’, 4’’-$CH_3$ with C-2’’, C-3’’ & C-4’’, and H-3’’ with 3’’-N-$CH_3$, as well as $^1$H-$^1$H COSY correlations of H-1’’/H-2’’/H-3’’ (Supplementary Figs. 3c–e). The linkage of these three fragments was inferred from HMBC correlations of H-4 with C-1’ and H-6 with C-1’’. The relative configuration of oxo-verdamicin (**6**) was assigned according to key NOESY correlations as shown in Supplementary Fig. 3f. Combined with the biosynthetic pathway, the structure of this compound was determined as shown in Supplementary Fig. 3g.

Considering that the conformation of the unsaturated sugar ring in oxo-verdamicin (**6**) in TFA were different from that in sisomicin (**7**), the $^1$H NMR spectra of oxo-verdamicin and sisomicin were acquired under acidic and basic conditions (Supplementary Fig. 3h). From pH 4.5 to pH 9.0, the conformation of unsaturated sugar ring in sisomicin (**7**) changed from $^1C_4$-like conformation to $^4C_1$-like conformation according to the H-3’ signal changes [pH 4.5: $\delta_H$ 2.34 (dt, 18.6, 3.4 Hz); pH 9.0: $\delta_H$ 2.02 (dd, 17.0, 9.1 Hz)]. This change was also observed in oxo-verdamicin (**6**): at pH 4.5 the unsaturated sugar ring exhibited a $^1C_4$-like conformation [$\delta_H$ 2.58 (dt, 20.8, 4.4 Hz)]; at pH 9.0, it displayed a mixture of both conformations (the ratio of $^1C_4$-like to $^4C_1$-like is 1:0.8); and finally at pH 12.0, it only exhibited the $^4C_1$-like conformation [$\delta_H$ 2.17 (dd, 18.6, 10.9 Hz)] (Supplementary Fig. 3h). The conformations of these compounds described here are shown in Supplementary Fig. 3i.

The molecular formula of verdamicin C2a (**8**) was deduced to be $C_{20}H_{39}N_5O_7$, according to the quasi-molecular ion peak at *m/z* 462.2887 $[M + H]^+$ (calc. for $C_{20}H_{40}N_5O_7^+$, 462.2922) in its ESI-HRMS spectrum (Supplementary Fig. 1j). Comparison of the $^1$H NMR spectra (Supplementary Fig. 6a) of verdamicin C2a (**8**) and oxo-verdamicin (**6**), showed that most of their signals were identical, except for two significant changes. The proton signal for H-4' at $\delta_H$ 6.43 in oxo-verdamicin (**6**) shifted to $\delta_H$ 5.22 in verdamicin C2a (**8**), and the singlet methyl signal for 6'-Me at $\delta_H$ 2.38 changed to a doublet at $\delta_H$ 1.47. These indicated that the C-6' ketone in oxo-verdamicin (**6**) was reduced and (from the molecular formula change) the oxygen at C-6' was replaced by an $NH_2$. The rest of the structure was confirmed by its $^1H$-$^1H$ COSY spectrum (Supplementary Figs. 6b). Thus, the planar structure of verdamicin C2a (**8**) was determined. Verdamicin C2 (**9**) had the same molecular formula as verdamicin C2 (**8**) according to the quasi-molecular ion peak at *m/z* 462.2888 $[M + H]^+$ (calc. for $C_{20}H_{40}N_5O_7^+$, 462.2922) in its ESI-HRMS spectrum (Supplementary Fig. 1k). Their $^1H$ NMR spectra (Supplementary Figs. 6a, d) are almost the same except for some minor differences, due to the different stereo-configuration of C-6'. All details other than the configuration at C-6' were confirmed through further analysis of the $^1H$ NMR data. Therefore, the structures were identified as verdamicin C2a (**8**) and verdamicin C2 (**9**), shown in Supplementary Fig. 6g.

**Comparison of the two verdamicin isomers (verdamicin 1 and verdamicin 2) isolated from Δ*genB4* with synthetic standards verdamicin C2a** (**8**) **and verdamicin C2** (**9**)[6] **by LC-MS.** LC-ESI-MS analysis was performed on a ThermoFinnigan LCQ connected to an Agilent HP 1100 HPLC system using a Phenomenex Prodigy C18 column (250×4.6 mm, 5 μ) at a flow rate of 0.3 mL/min, with a mobile phase of (A) 0.2% TFA in $H_2O$ and (B) 100% $CH_3CN$; the gradient used was: 0-15 min 2% B to 5% B, 15-22 min 5% B to 95% B, 22-33 min 95% B. MS/MS analyses were carried out in the positive ionization mode with 35% relative collision energy.

**Protein expression and purification for crystallization assays.** The *genB3* and *genB4* gene from *M. echinospora* ATCC15835 were cloned into plasmid pET28a(+) and expressed in *E. coli* BL21(DE3) cells (Novagen) as described above. Briefly, cells were harvested by centrifugation and resuspended in lysis buffer (50 mM HEPES (pH

8.0), 150 mM NaCl in the presence of lysozyme (1 mg/mL), DNase I (1 mg/mL) and phenylmethanesulfonyl fluoride (10 mM). Cells were disrupted by sonication (5 min with a 50% duty cycle), and the lysate was clarified by centrifugation (15000 g, 4°C, 1 h). The supernatant was passed through a 5 mL IMAC column (GE Healthcare) charged with nickel and previously equilibrated with 50 mM HEPES (pH 8.0) and 150 mM NaCl. The proteins were eluted using a linear gradient of imidazole (up to 500 mM) in a buffer of 50 mM HEPES (pH 8.0) and 150 mM NaCl on an ÅKTA Purifier (GE Healthcare). Fractions containing the proteins were concentrated using Amicon-Ultra Centrifugal Filters (Millipore) and further purified by gel filtration on a 16/60 Superdex 200 column (GE Healthcare) equilibrated in 50 mM HEPES (pH 8.0) and 150 mM NaCl. Fractions containing GenB3 or GenB4 were concentrated to 15−20 mg/mL, and the enzyme was stored at −80 °C until use. The N-terminal His-tag was retained in the subsequent experiments.

**GenB3 and GenB4 crystallization.** Crystallization trials were carried out using the sitting-drop method. The initial trials were performed using an Oryx robot (Douglas Instruments) at the Laboratory of Applied Structural Biology at São Paulo-Brazil using the sitting drop method in 96 well plates. GenB3 and GenB4 at 10 mg/mL were incubated in the presence of 5 mM PLP or alternatively in the presence of 5 mM PLP and 10 mM sisomicin (**7**) and submitted to crystallization screening kits from Jene bioscience (Classic I-IV, Basic I-IV, JCSG++ I-IV, Pi-minimal) and Hampton Research (Index). The drops were comprised of 0.2 μL of protein solution and 0.2 μL of crystallization solution and the plates were stored at 18 °C. The best conditions were optimized manually by hanging drop vapor diffusion method using 24-well Linbro plates. The best and most reproducible crystallization conditions for GenB3 were 100 mM sodium malonate, pH 4.0 and 12% (v/v) PEG 3350 and generally, the crystals appeared after 7 days. Crystals of GenB3 in presence of sisomicin (**7**) were not obtained. GenB4-PLP internal aldimine or the external aldimine complexes were obtained in the presence of PLP or in the presence of both PLP and sisomicin (**7**), respectively, in 100 mM DL-malic acid, pH 7.0 and 20% (v/v) PEG 3350.

**Data Collection and Processing and Structure Determination.** Data collection of GenB3-PLP was carried out at MX2 National Synchrotron Light Laboratory (Campinas-Brazil) and GenB4-PLP and GenB4-PLP-**7** data were collected at PETRA III, Hamburg, Germany, P13 beamline. The data was processed using XDS[7] and scaled using AIMLESS[8] from CCP4i[9]. The structure of GenB4 was determined through the MrBump server[10], which uses an algorithm that through experimental structure and target sequence factors, searches in the servers chains, multimeters and domains to be used in molecular substitution in combination with the Arp/warp program[11] that constructs the model of the protein from the generated results of the molecular substitution. The structure of GenB4-PLP-**12** external aldimine and GenB3-PLP internal aldimine were determined by molecular substitution using Phaser[12] from Phenix crystallographic suite[13] using the GenB4:PLP structure as a search model. The structures were refined using Phenix.refine[14]. Manual building, visual inspection, and analysis were carried out using COOT[15] and further analysis and figure preparation used the PyMOL Molecular Graphics System, Version 1.8 (Schrodinger, LLC). The quality of the structure was checked using the program Molprobity[16].

**Supplementary Table 1 | $^{1}H$ (600 MHz) and $^{13}C$ (150 MHz) NMR results of oxo-verdamicin (6) in $D_2O$.**

| No. | $^{13}C$ | $^{1}H$ |
|---|---|---|
| 1 | 51.1, d | 3.57 (m, 1H) |
| 2 | 28.9, t | 2.55 (dt, 12.6, 4.3 Hz, 1H); |
| | | 1.94 (q, 12.6 Hz, 1H) |
| 3 | 49.5, d | 3.54 (m, 1H) |
| 4 | 81.5, d | 4.03 (dd, 10.1, 9.2 Hz, 1H) |
| 5 | 74.7, d | 3.79 (t, 9.2 Hz, 1H) |
| 6 | 84.5, d | 3.76 (t, 9.2 Hz, 1H) |
| 1′ | 98.5, d | 5.62 (s, 1H) |
| 2′ | 47.1, d | 3.99 (m, 1H) |
| 3′ | 25.4, t | 2.93 (ddd, 20.0, 5.7, 4.3 Hz, 1H); |
| | | 2.62 (dt, 20.0, 4.3 Hz, 1H) |
| 4′ | 115.1, d | 6.43 (t, 4.3 Hz, 1H) |
| 5′ | 148.1, s | — |
| 6′ | 197.3, s | — |
| 6′-$CH_3$ | 25.9, q | 2.38 (s, 3H) |
| 1″ | 102.6, d | 5.09 (d, 3.7 Hz, 1H) |
| 2″ | 67.6, d | 4.23 (dd, 10.9, 3.7 Hz, 1H) |
| 3″ | 64.7, d | 3.47 (d, 10.9 Hz, 1H) |
| 3″-N-$CH_3$ | 35.8, q | 2.92 (s, 3H) |
| 4″ | 71.2, s | — |
| 4″-$CH_3$ | 22.2, q | 1.35 (s, 3H) |
| 5″ | 69.0, t | 3.99 (d, 12.9 Hz, 1H); |
| | | 3.50 (d, 12.9 Hz, 1H) |

**Supplementary Table 2 | $^1$H NMR results (600 MHz) of verdamicin C2a (8) and verdamicin C2 (9) in $D_2O$**

| No. | $^1$H | |
|---|---|---|
| | verdamicin C2a (8) | verdamicin C2 (9) |
| 1 | 3.57 (m, 1H) | 3.57 (m, 1H) |
| 2 | 2.56 (dt, 12.6, 4.0 Hz, 1H);<br>1.96 (q, 12.6 Hz, 1H) | 2.56 (dt, 12.6, 4.3 Hz, 1H);<br>1.96 (q, 12.6 Hz, 1H) |
| 3 | 3.54 (m, 1H) | 3.54 (m, 1H) |
| 4 | 4.04 (t, 9.8 Hz, 1H) | 4.02 (m, 1H) |
| 5 | 3.81 (t, 9.4 Hz, 1H) | 3.81 (t, 9.4 Hz, 1H) |
| 6 | 3.75 (t, 9.4 Hz, 1H) | 3.75 (t, 9.4 Hz, 1H) |
| 1′ | 5.71 (s, 1H) | 5.66 (s, 1H) |
| 2′ | 3.91 (t, 5.1 Hz, 1H) | 3.94 (m, 1H) |
| 3′ | 2.70 (dt, 18.4, 4.3 Hz, 1H);<br>2.42 (dt, 18.4, 4.3 Hz, 1H) | 2.72 (ddd, 18.8, 5.7, 3.7 Hz, 1H);<br>2.39 (dt, 18.8, 3.7 Hz, 1H) |
| 4′ | 5.22 (t, 3.5 Hz, 1H) | 5.18 (t, 3.5 Hz, 1H) |
| 5′ | — | — |
| 6′ | 3.99 (m, 1H) | 4.04 (q, 6.8 Hz, 1H) |
| 6′-$CH_3$ | 1.47 (d, 6.9 Hz, 3H) | 1.47 (d, 6.8 Hz, 3H) |
| 1″ | 5.08 (d, 3.6 Hz, 1H) | 5.08 (d, 3.6 Hz, 1H) |
| 2″ | 4.23 (dd, 10.9, 3.6 Hz, 1H) | 4.23 (dd, 10.9, 3.6 Hz, 1H) |
| 3″ | 3.47 (d, 10.9 Hz, 1H) | 3.47 (d, 10.9 Hz, 1H) |
| 3″-N-$CH_3$ | 2.92 (s, 3H) | 2.92 (s, 3H) |
| 4″ | — | — |
| 4″-$CH_3$ | 1.35 (s, 3H) | 1.35 (s, 3H) |
| 5″ | 3.99 (d, 12.9 Hz, 1H);<br>3.50 (d, 12.9 Hz, 1H) | 3.99 (d, 12.9 Hz, 1H);<br>3.50 (d, 12.9 Hz, 1H) |

**Supplementary Table 3 | X-ray data analysis of GenB3 and GenB4 structures.**

| | GenB3-PLP | GenB4-PLP | GenB4-PLP-12 |
|---|---|---|---|
| PDB entry | 7LM0 | 7LLE | 7LLD |
| X-ray Source | PETRA III-P13/ Germany | LNLS-MX2 /Brazil | PETRA III-P13/ Germany |
| Wavelength (Å) | 0.97 | 1.45 | 0.97 |
| Resolution range (Å) | 31.3 - 2.1 (2.2 - 2.1) | 46.0 - 1.7 (1.8 - 1.7) | 40.0 - 1.4 (1.4 - 1.4) |
| Space group | P 1 21 1 | P 21 21 21 | P 21 21 21 |
| Unit cell (Å) | 59.5 156.7 63.9 90 115.197 90 | 71 74.2 181 90 90 90 | 71.4 75 186 90 90 90 |
| Total reflections | 214698 | 209969 (20513) | 387282 (37415) |
| Unique reflections | 59024 (5610) | 105005 (10268) | 195482 (19105) |
| Multiplicity | 1.9 (1.8) | 2.0 (2.0) | 2.0 (2.0) |
| Completeness (%) | 94.83 (90.54) | 99.79 (98.77) | 99.54 (98.50) |
| Mean I/Sigma(I) | 12.72 (1.72) | 15.9 (1.85) | 12.85 (1.62) |
| Wilson B-factor (Å) | 36.14 | 18.38 | 16.83 |
| R-merge | 0.031 (0.38) | 0.03 (0.4) | 0.03 (0.5) |
| R-pim | 0.021 (0.54) | 0.03 (0.4) | 0.03 (0.5) |
| CC1/2 | 0.998 (0.77) | 0.99 (0.7) | 0.99 (0.64) |
| Reflections used in refinement | 59018 (5610) | 104930 (10268) | 195052 (19089) |
| R-work | 0.161 (0.211) | 0.154 (0.244) | 0.145 (0.227) |
| R-free | 0.196 (0.286) | 0.185 (0.278) | 0.188 (0.281) |
| Number of non-hydrogen atoms | 7583 | 8197 | 8479 |
| Macromolecules | 6913 | 6963 | 7063 |
| Ligands | 37 | 30 | 109 |
| Solvent | 633 | 1204 | 1307 |

| | | | |
|---|---|---|---|
| Protein residues | 896 | 891 | 892 |
| RMS (bonds) | 0.008 | 0.006 | 0.006 |
| RMS (angles) | 0.96 | 0.84 | 1.08 |
| Ramachandran Favored (%) | 96.86 | 97.18 | 97.29 |
| Ramachandran Allowed (%) | 2.92 | 2.59 | 2.48 |
| Ramachandran Outliers (%) | 0.22 | 0.23 | 0.23 |
| Rotamer outliers (%) | 0.14 | 0.28 | 0.41 |
| Clashscore | 4.02 | 2.97 | 5.60 |
| Average B-factor | 38.79 | 22.81 | 24.78 |
| Macromolecules | 38.18 | 20.29 | 21.33 |
| Ligands | 29.67 | 15.04 | 41.18 |
| Solvent | 46.00 | 37.58 | 42.06 |
| Number of TLS Groups | 1 | 1 | 0 |

Statistics for the highest-resolution shell are shown in parentheses.

**Supplementary Table 4 | Oligonucleotide primers used for in-frame deletions and complementations in this study.**

| Primer | Oligonucleotide sequence (5’ to 3’) |
|---|---|
| B3-in-K-L-1 | CGT**CATATG**GCAACACCACGTCG (*Nde*I) |
| B3-in-K-L-2 | GAC**GAGCTC**ATCGAGAAGGTGGTC (*Sac*I) |
| B3-in-K-R-1 | CGA**GAGCTC**GGTCCCGATGTCGTAG (*Sac*I) |
| B3-in-K-R-2 | CAGACG**AAGCTT**AACGCGGCACCGG (*Hin*dIII) |
| genP-L1 | CAG**CATATGC**TGGATGCGGTGGTC (*Nde*I) |
| genP-L2 | GAC**GAATTC**CTCTGAGCTGACCCGG (*Eco*RI) |
| genP-R1 | CGG**GAATTC**CTTGTCGCCCCAGCC (*Eco*RI) |
| genP-R2 | GGG**AAGCTT**GCGGGAAAGTCGACCA (*Hin*dIII) |
| genB3-in-K -CK1 | CCTCCTTGGTCGGGTTGA |
| genB3-in-K -CK2 | CGTCGCGTTACGGAAAGT |
| genB4-CK1 | TGACTTCTGCCTCGACAACG |
| genB4-CK2 | AAGCTCTACCTGGAGACCTTCC |
| genK-CK1 | CGGGCGAACCTTCGGGATA |
| genK-CK2 | CCGTCAGCGTTGGCAATAA |
| genP-CP1 | ATGACGGTAGCCGAGGATG |
| genP-CP2 | GCGTTGACGGCGTTCC |
| 77-B3-P)-B2-gmrA | CATCAGCGAAACCTCCGGTCAGAGAAATTCGTCCAGCA |
| 77-B3-P-(B2-gmrA | TGCTGGACGAATTTCTCTGACCGGAGGTTTCGCTGATG |
| 77-B3-P-B4)-gmrA | TCCTCCGAAAGATCCTTCAGTTCTGTGCCGGGAA |
| 77-(B4 | TTGGTAGGATCCACATATGAACTACCGTGAGTTGATCGAG |
| 77-B4-B3-(gmrA | AGGATCTTTCGGAGGACT |
| 77-B4-B3)-gmrA | TCCTCCGAAAGATCCTTCAGTTCTGGGCGGGGA |
| 77-B4)-B3-gmrA | GTCCTGGCTACCTTCTCTCAGTTCTGTGCCGG |
| 77-B4-(B3-gmrA | GAGAAGGTAGCCAGGACATGGATTCTGCCAACT |
| P)-gmrA | GAAAGATCCTTCAGAGAAATTCGTCCAGCAGTTGG |
| P-(gmrA | GAATTTCTCTGAAGGATCTTTCGGAGGACTCGATG |
| 77-(B3PB4-B2-gmrA | TTGGTAGGATCCACATATGGATTCTGCCAACTTG |

| | |
|---|---|
| 77-B3PB4)-B2-gmrA | CGCCCCTCGGTCAGTTCTGTGCCGGGAAGA |
| 77-B3PB4-(B2-gmrA | ACAGAACTGACCGAGGGGCGCAGAGGACAG |
| 77-B3PB4-B2)-gmrA | ACCGGCAGCGTCAGAGCTGAGCGGTCACGT |
| 77-B3PB4-B2-(gmrA | TCAGCTCTGACGCTGCCGGTTCAGGATCTT |
| 77-B3PB4-B2-gmrA) | GACATGATTACGAATTCCTATTTCTGAATGACGTAA |
| a-77-(B2 | TTGGTAGGATCCACATATGATTATTGCCAACGCT |
| a-B2)-gmrA | TCCTCCGAAAGATCCTTCAGAGCTGAGCGGTC |
| a-77-B4)-gmrA | GGGGAGGACGTGAAATTCTGCCCGGGTCAG |
| a-77-B4-(gmrA | CTGACCCGGGCAGAATTTCACGTCCTCCCCA |
| a-77-B4-gmrA) | ACCCGGGCAGAATTCAAGCTATTTCTGAATGACG |
| a-B2-(gmrA | TCAGCTCTGACGAATTCGGTAATACTCCTGC |
| a-B2-gmrA)-B4 | GCCATTCACGTCTCCAAGCTATTTCTGAATGACG |
| a-B2-gmrA)-B3 | GACCTGTGGTCGGCGAAGCTATTTCTGAATGACG |
| a-B2-gmrA-(B4 | GGAGACGTGAATGGCGTG |
| a-B2-gmrA-B4)-77 | ACATGATTACGAATTGGTCAGTTCTGTGCCGG |
| a-B2-gmrA-(B3-77 | CGCCGACCACAGGTCGAGC |
| a-B2-gmrA-B3)-77 | ACATGATTACGAATTTCAGTTCTGGGCGGGGA |
| EP-genB2-CK1 | ATGATTATTGCCAACGCTGACGGTTG |
| EP-genB2-CK2 | TCAGAGCTGAGCGGTCACGTACTCC |
| EP-genB3-CK1 | TCAGTTCTGGGCGGGGATGAGAACC |
| EP-genB3-CK2 | ATGGATTCTGCCAACTTGACGAACC |
| EP-genB4-CK1 | TCAGTTCTGTGCCGGGAAGAGGACC |
| EP-genB4-CK2 | ATGAACTACCGTGAGTTGATCGAGCGG |
| EP-genP-CK1 | TCAGAGAAATTCGTCCAGCAGTTGG |
| EP-genP-CK2 | ATGGTTGCAGCACCGATACCGGTGG |
| EP-gmrA-CK1 | CTATTTCTGAATGACGTAAATCAG |
| EP-gmrA-CK2 | ATGACGACATCTGCGCCTGAGGACC |

**Supplementary Table 5 | Plasmids used in this study.**

| Plasmid | Description | Reference |
|---|---|---|
| pUC18 | Sub-cloning vector | 17 |
| pYH7 | *Streptomyces-E. coil* shuttle vector | 5 |
| pWHU77 | pIB139 derived integrative vector with $Tsr^R$ | 2 |
| pWHU166 | pYH7 derived recombinant plasmid used for in-frame deletion of *genP* | This study |
| pWHU2740 | pYH7 derived recombinant plasmid used for in-frame deletion of *genB3* in ΔgenK | This study |
| pWHU1 | pYH7 derived recombinant plasmid used for in-frame deletion of *genK* in ΔgenP | 2 |
| pWHU3 | pYH7 derived recombinant plasmid used for in-frame deletion of *genB4* in ΔgenK | 2 |
| pWHU90 | Recombinant plasmid used for complementation of *gmrA*, which is inserted into vector pWHU77 under the control of the P*ermE** promoter. | 4 |
| pWHUB3inK | pYH7 derived recombinant plasmid used for in-frame deletion of *genB3* in ΔgenK | This study |
| pWHU81 | Recombinant plasmid used for complementation of *genB3* and *gmrA*, which are inserted into pWHU77, under the control of the same P*ermE** promoter. The fragment of *genB3* is achieved with primers 77-(B3PB4-B2-gmrA and 77-B4-B3)-gmrA. The fragment of *gmrA* is achieved with primer 77-B4-B3-(gmrA and a-77-B4-gmrA) | This study |
| pWHU82 | Recombinant plasmid used for complementation of *genB4* and *gmrA*, which are inserted into pWHU77, under the control of the same P*ermE** promoter. The fragment of *genB4* is achieved | This study |

| | | |
|---|---|---|
| | with primers 77-(B4 and a-77-B4)-gmrA. The fragment of *gmrA* is achieved with primer a-77-B4-(gmrA and a-77-B4-gmrA) | |
| pWHU83 | Recombinant plasmid used for complementation of *genB2* and *gmrA*, which are inserted into pWHU77, under the control of the same P*ermE** promoter. The fragment of *genB2* is achieved with primers a-77-(B2 and a-B2)-gmrA. The fragment of *gmrA* is achieved with primer a-B2-(gmrA and a-77-B4-gmrA) | This study |
| pWHU84 | Recombinant plasmid used for complementation of *genB2*, *gmrA* and *genB4*, which are inserted into pWHU77, under the control of the same P*ermE** promoter. The fragment of *genB2* is achieved with primers a-77-(B2 and a-B2)-gmrA. The fragment of *gmrA* is achieved with a-B2-(gmrA and a-B2-gmrA)-B4. The fragment of *genB4* is achieved with a-B2-gmrA-(B4 and a-B2-gmrA-B4)-77 | This study |
| pWHU86 | Recombinant plasmid used for complementation of *genB4*, *genB3* and *gmrA*, which are inserted into pWHU77, under the control of the same P*ermE** promoter. The fragment of *genB4* is achieved with primers 77-(B4 and 77-B4)-B3-gmrA. The fragment of *genB3* is achieved with 77-B4-(B3-gmrA and 77-B4-B3)-gmrA. The fragment of *gmrA* is achieved with 77-B4-B3-(gmrA and a-77-B4-gmrA) | This study |
| pWHU87 | Recombinant plasmid used for complementation | This study |

| | | |
|---|---|---|
| | of *genB3*, *genP* and *gmrA*, which are inserted into pWHU77, under the control of the same P*ermE** promoter. The fragment of *genB3*-*genP* is achieved with primers 77-(B3PB4-B2-gmrA and P)-gmrA. The fragment of *gmrA* is achieved with P-(gmrA and a-77-B4-gmrA). | |
| pWHU88 | Recombinant plasmid used for complementation of *genB3*, *genP*, *genB4* and *gmrA*, which are inserted into pWHU77, under the control of the same P*ermE** promoter. The fragment of *genB3*-*genP*-*genB4* is achieved with primers 77-(B3PB4-B2-gmrA and a-77-B4)-gmrA. The fragment of *gmrA* is achieved with primer a-77-B4-(gmrA and a-77-B4-gmrA) | This study |

**Supplementary Table 6 | In-frame deletion mutants and complementation strains used in this study.**

| Strain | Parent strain | Plasmid | Reference |
|---|---|---|---|
| ΔBN | wild-type | pWHU39 | 4 |
| Δ*genK* | wild-type | pWHU1 | 2 |
| Δ*genP*Δ*genK* | ΔgenP | pWHU1 | This study |
| Δ*genK*Δ*genB3* | ΔgenK | pWHU2740 | This study |
| Δ*genB4*Δ*genK* | ΔgenK | pWHU3 | This study |
| Δ*genP* | wild-type | pWHU166 | This study |
| Δ*genB3* | wild-type | pWHU5 | 2 |
| Δ*genB4* | wild-type | pWHU3 | 2 |
| ΔBN::*gmrA* | ΔBN | pWHU90 | 4 |
| ΔBN::*genB2-gmrA* | ΔBN | pWHU83 | This study |
| ΔBN::*genB3-gmrA* | ΔBN | pWHU81 | This study |
| ΔBN::*genB4-gmrA* | ΔBN | pWHU82 | This study |
| ΔBN::*genB2-gmrA-genB4* | ΔBN | pWHU84 | This study |
| ΔBN::*genB4-genB3-gmrA* | ΔBN | pWHU86 | This study |
| ΔBN::*genB3-genP-gmrA* | ΔBN | pWHU87 | This study |
| ΔBN::*genB3-genP-genB4-gmrA* | ΔBN | pWHU88 | This study |

**Supplementary Table 7 | Oligonucleotide primers used for site-directed mutations in this study.**

| Protein | Mutation | Primer | Primer sequence (5' to 3') |
|---|---|---|---|
| GenB3 | S57D | S57D_F | TCACCGCCTCC***GA***CGGGACGATCAT |
| | | S57D_R | ATGATCGTCCCG***T***CGGAGGCGGTGA |
| | S116T | S116T_F | ACAAGACCGGC***A***CCGAGGGTACGGC |
| | | S116T_R | GCCGTACCCTCGG***T***GCCGGTCTTGT |
| GenB4 | D52S | D52S_F | CCTGACCGGCGCC***AGC***GCCGCCGTCATCCTCGGCTA |
| | | D52S_R | TAGCCGAGGATGACGGCGGC***GCT***GGCGCCGGTCAGG |
| | T111S | T111S_F | ACAAAACTGGTA***G***CGAGGGCACGGC |
| | | T111S_R | GCCGTGCCCTCG***C***TACCAGTTTTGT |

Mutated nucleotides are in bold face and italicized.

**Supplementary Fig. 1 | LC-ESI-HRMS and MS/MS analysis of gentamicins and its intermediates.**

**a**, LC-ESI-HRMS and MS/MS analysis of JI-20A (**1**).

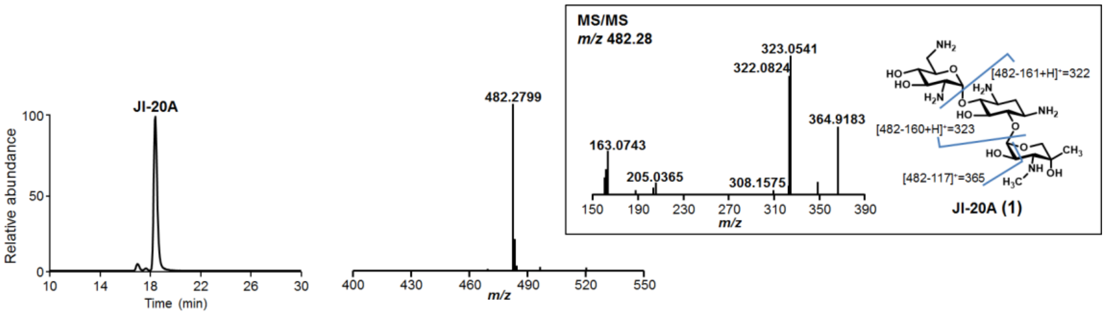


**b**, LC-ESI-HRMS and MS/MS analysis of JI-20Ba (**2**).

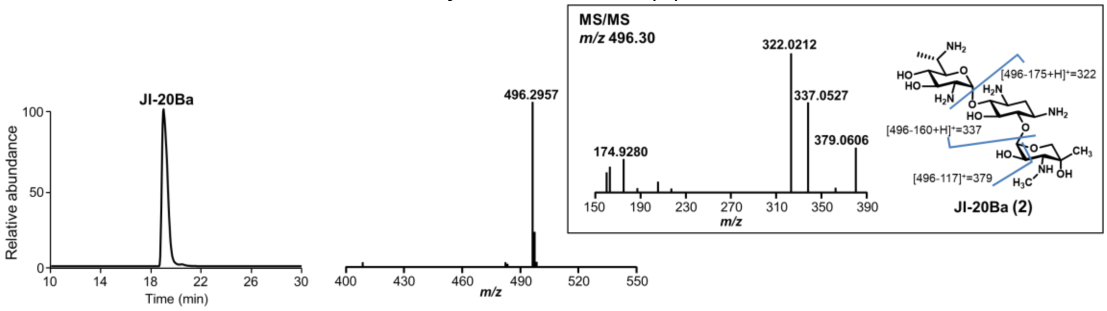


**c**, LC-ESI-HRMS and MS/MS analysis of JI-20A-P (**3**).

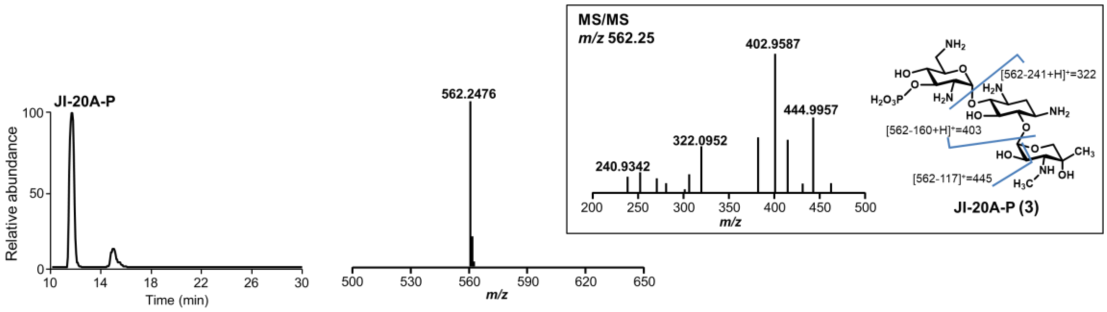


**d**, LC-ESI-HRMS and MS/MS analysis of JI-20Ba-P (**4**).

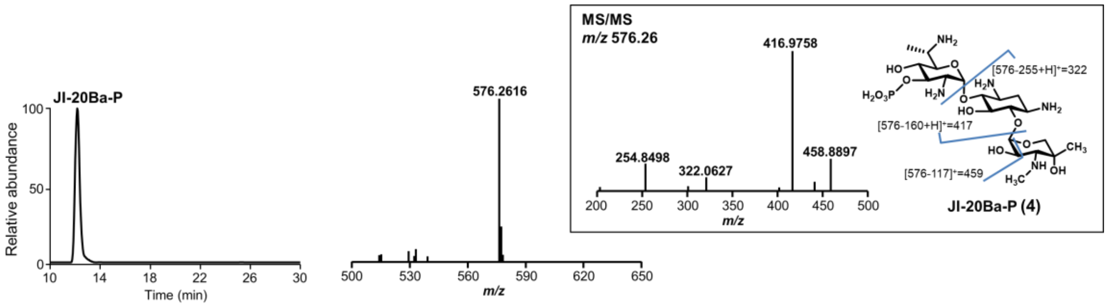

**e**, LC-ESI-HRMS and MS/MS analysis of oxo-sisomicin (**5**).

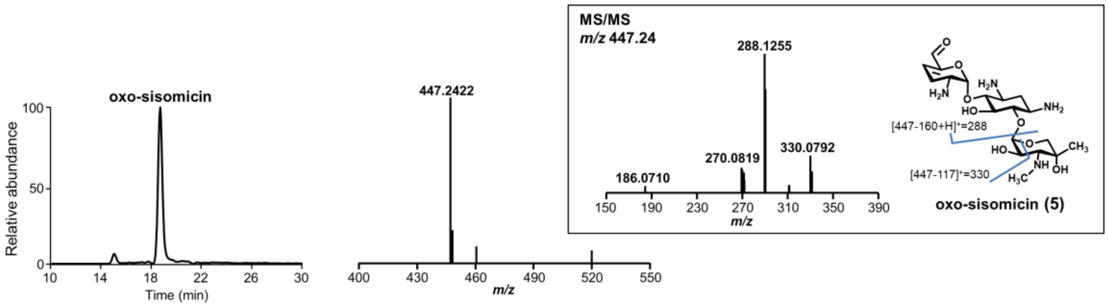


**f**, LC-ESI-HRMS and MS/MS analysis of oxo-verdamicin (**6**).

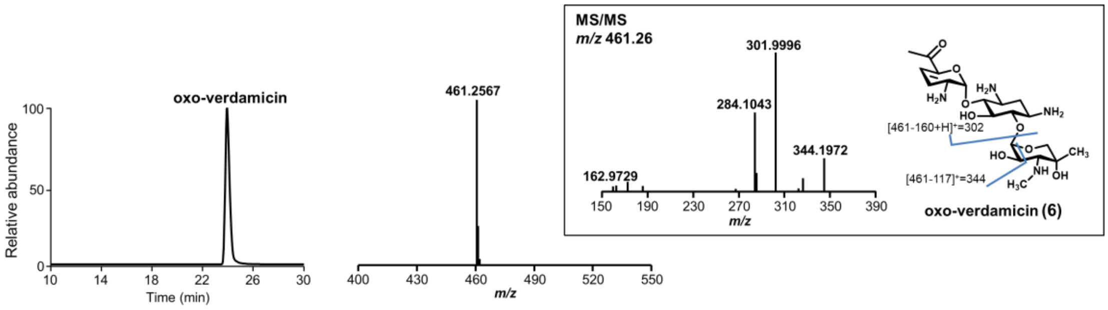


**g**, LC-ESI-HRMS and MS/MS analysis of sisomicin (**7**).

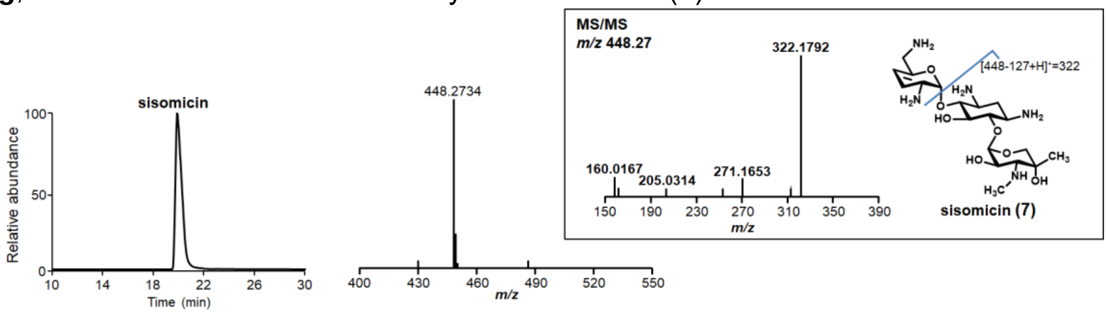


**h**, LC-ESI-HRMS and MS/MS analysis of C1a (**12**).

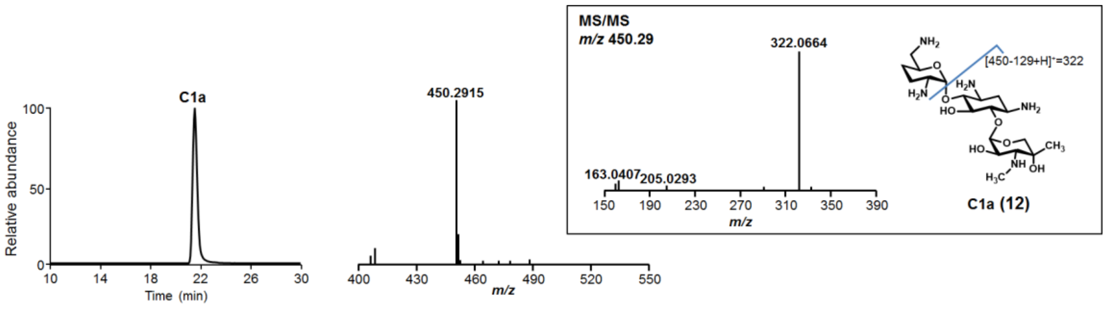

**i**, LC-ESI-HRMS and MS/MS analysis of C2b (**14**).

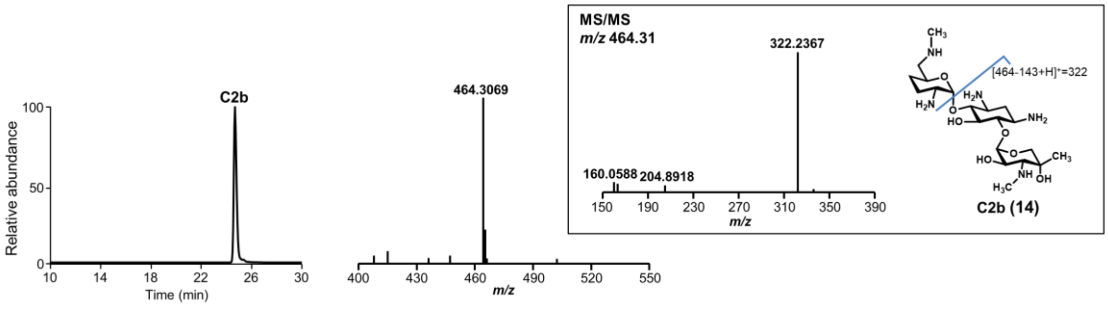


**j**, LC-ESI-HRMS and MS/MS analysis of verdamicin C2a (**8**).

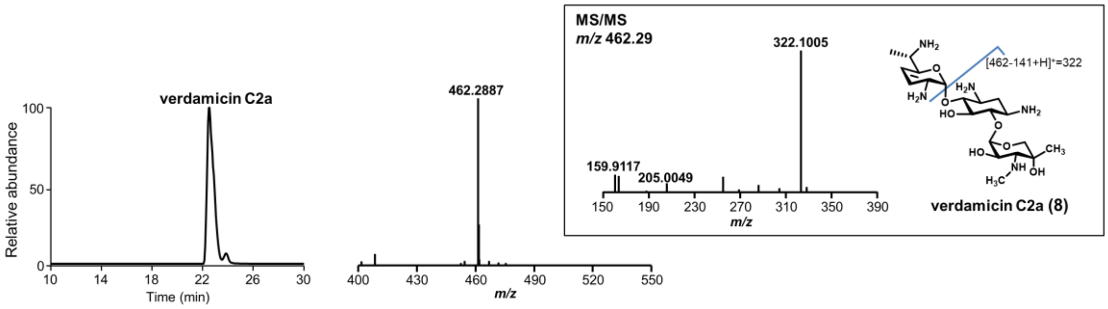


**k**, LC-ESI-HRMS and MS/MS analysis of verdamicin C2 (**9**).

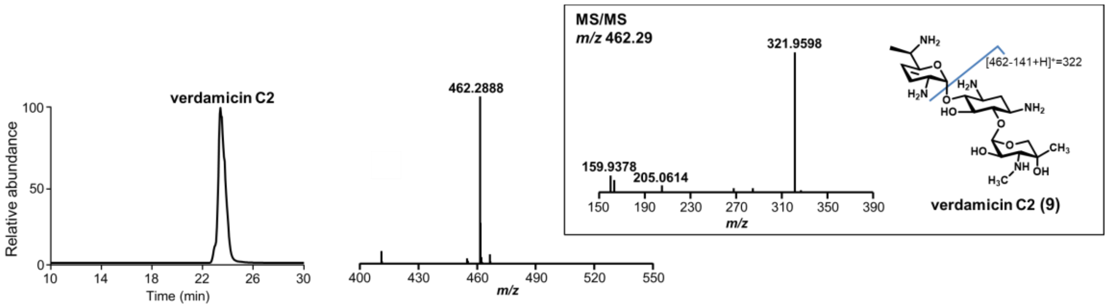


**l**, LC-ESI-HRMS and MS/MS analysis of oxo-C1a (**10**).

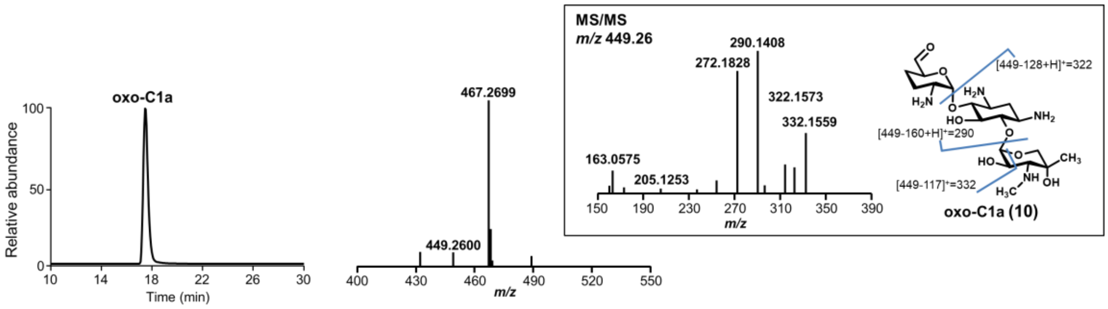

**m**, LC-ESI-HRMS and MS/MS analysis of oxo-C2a (**11**).

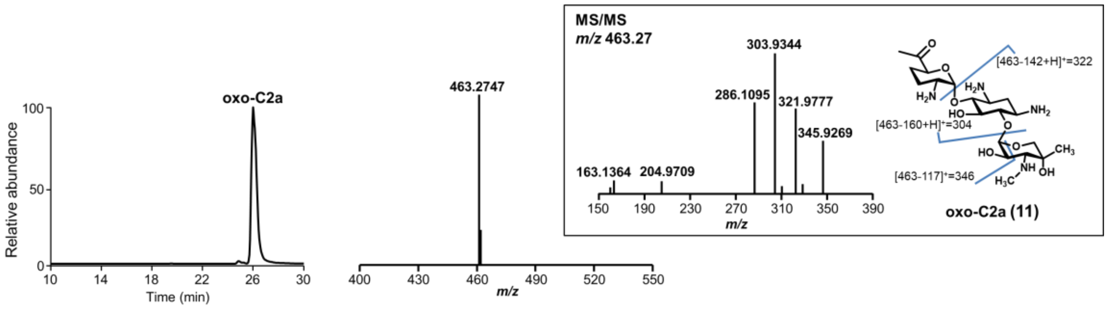


**n**, LC-ESI-HRMS and MS/MS analysis of C2a (**13**).

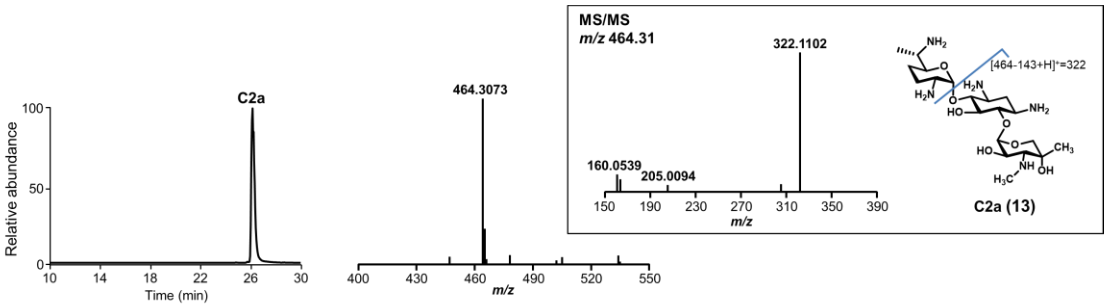


**o**, LC-ESI-HRMS and MS/MS analysis of C2 (**15**).

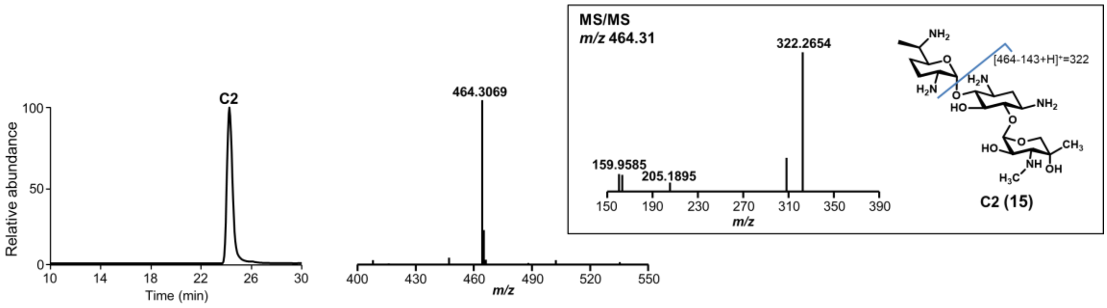


**p**, LC-ESI-HRMS and MS/MS analysis of C1 (**16**).

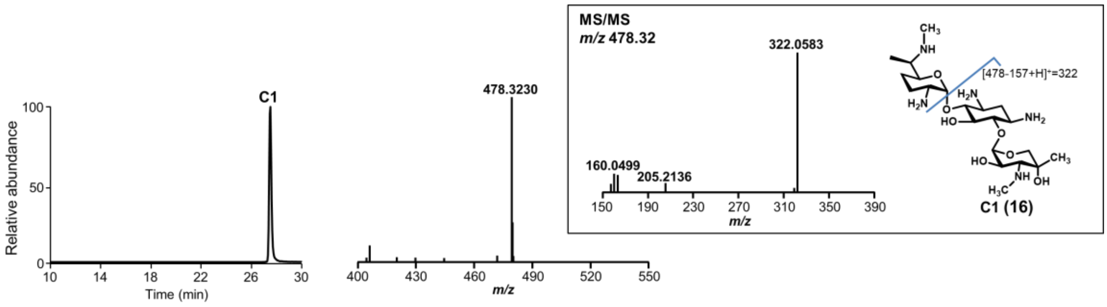

**q**, LC-ESI-HRMS and MS/MS analysis of JI-20Bb.

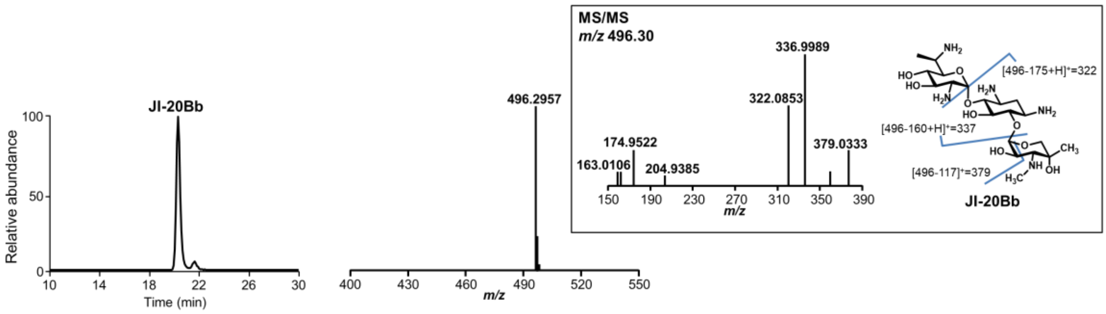


**r**, LC-ESI-HRMS and MS/MS analysis of JI-20Bb-P.

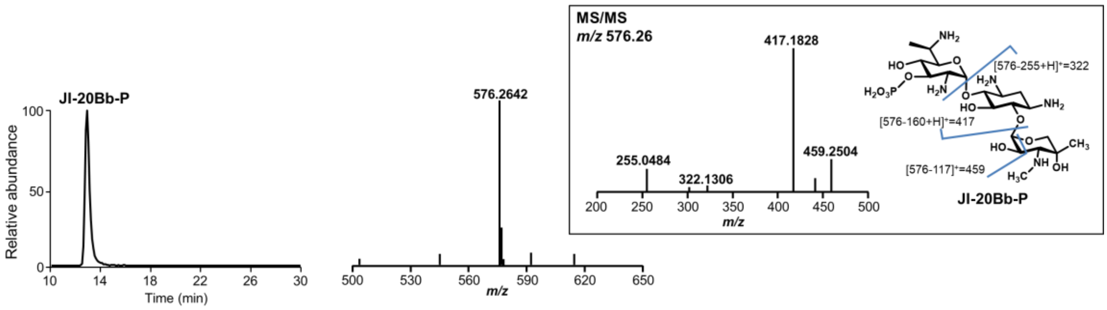

**Supplementary Fig. 2 | Confirmation of in-frame deletion mutants.**

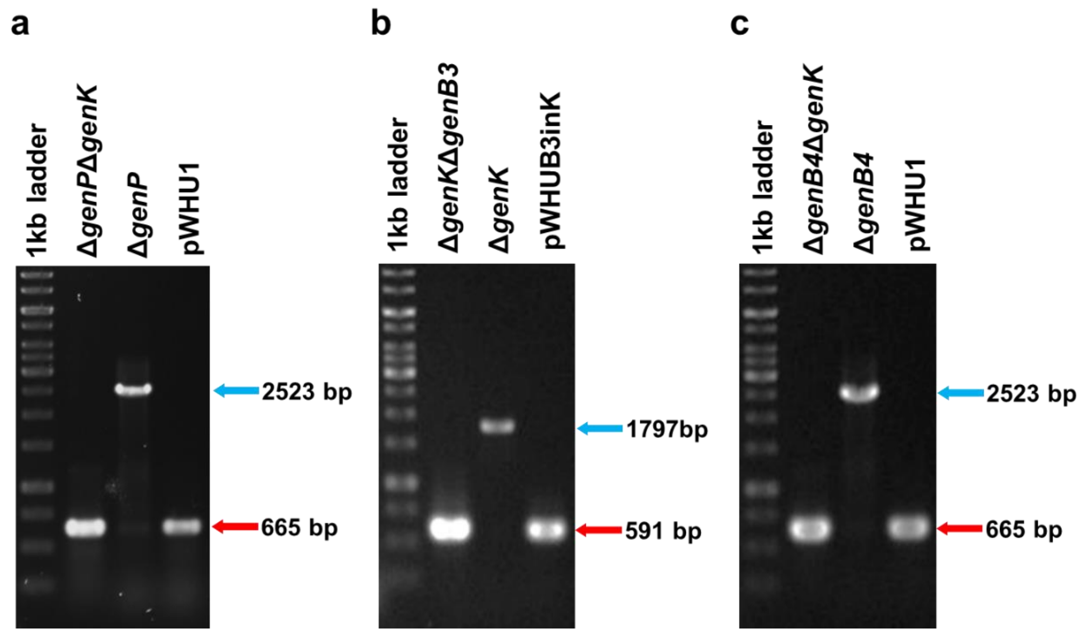


Confirmation of in-frame deletion mutants of (**a**) Δ*genP*Δ*genK*, (**b**) Δ*genK*Δ*genB3* and (**c**) Δ*genB4*Δ*genK* by PCR. The primers used are listed in Supplementary Table 1. The blue arrows indicate the expected size of PCR fragments of parent strain, and the red arrows indicate those of mutants.

**Supplementary Fig. 3 | NMR data and analysis of oxo-verdamicin (6) in $CF_3CO_2D$.**

**a**, $^{1}$H-NMR spectrum (600 MHz) of keto-verda (**6**).

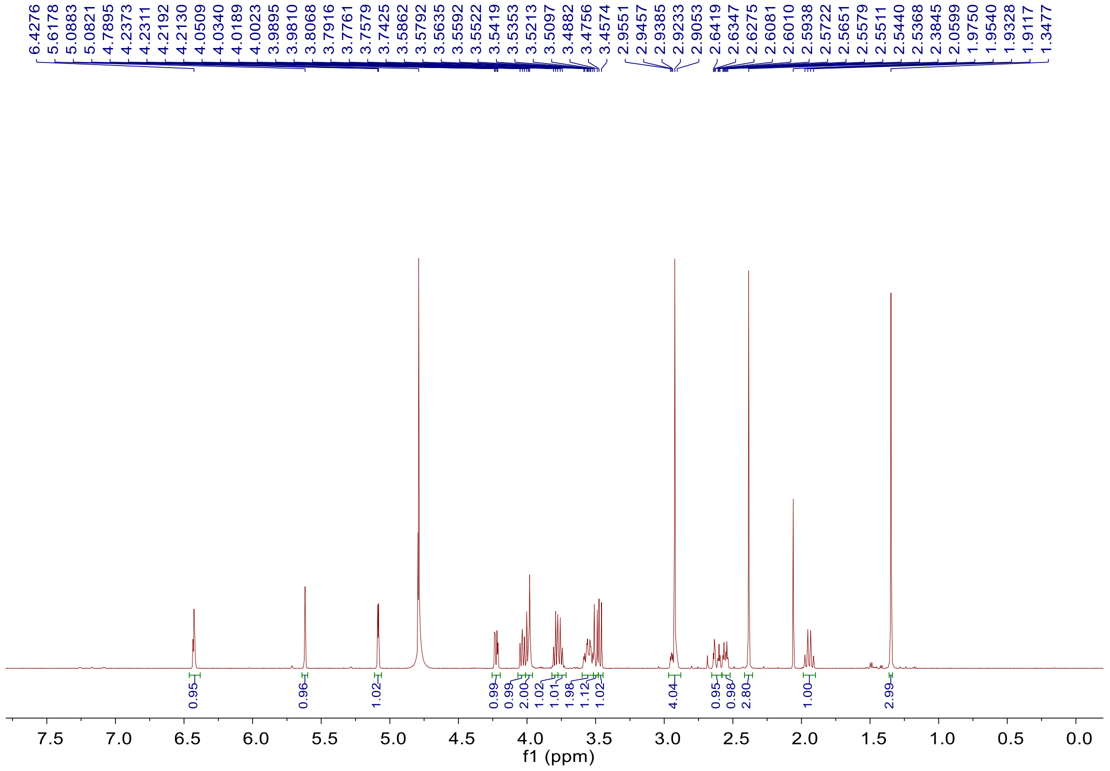


**b**, $^{13}$C-NMR and DEPT spectrum (150 MHz) of oxo-verdamicin (**6**).

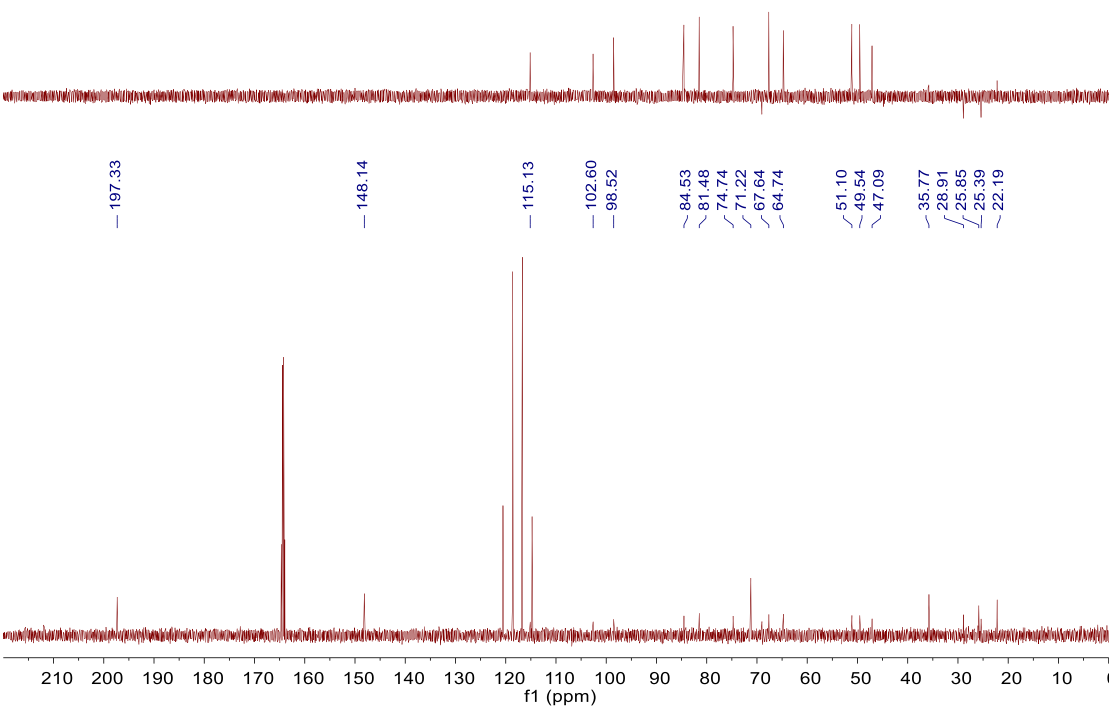

**c**, $^{1}H$-$^{1}H$ COSY Spectrum (600 MHz) of oxo-verdamicin (**6**).

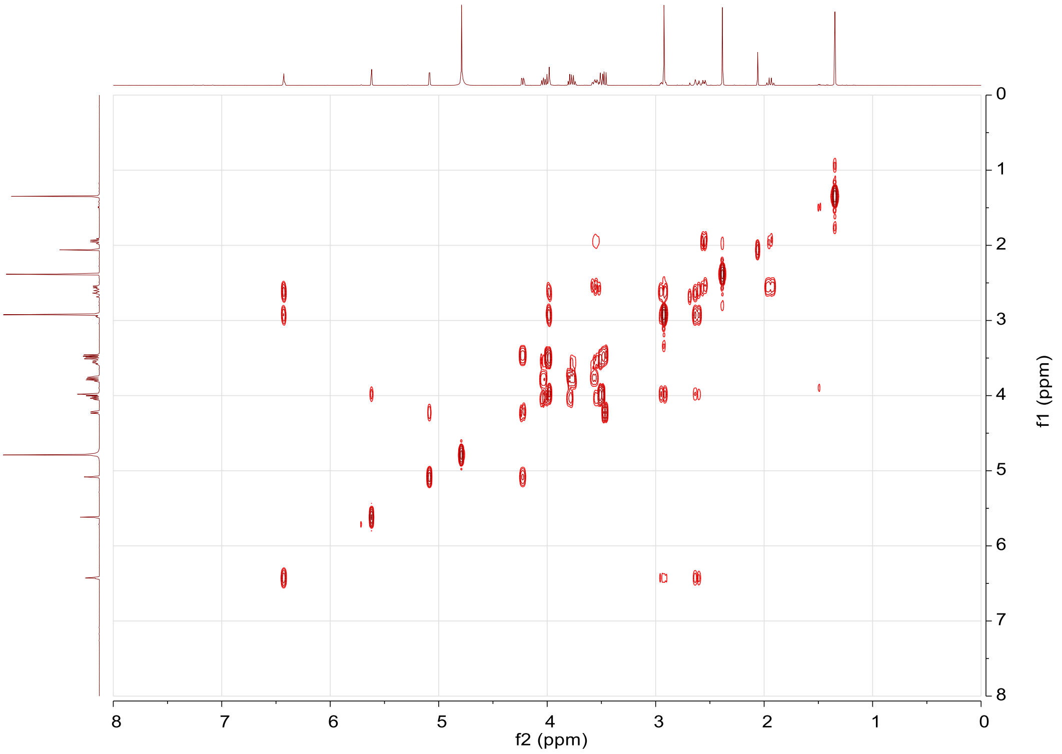


**d**, HSQC spectrum (600 MHz) of oxo-verdamicin (**6**).

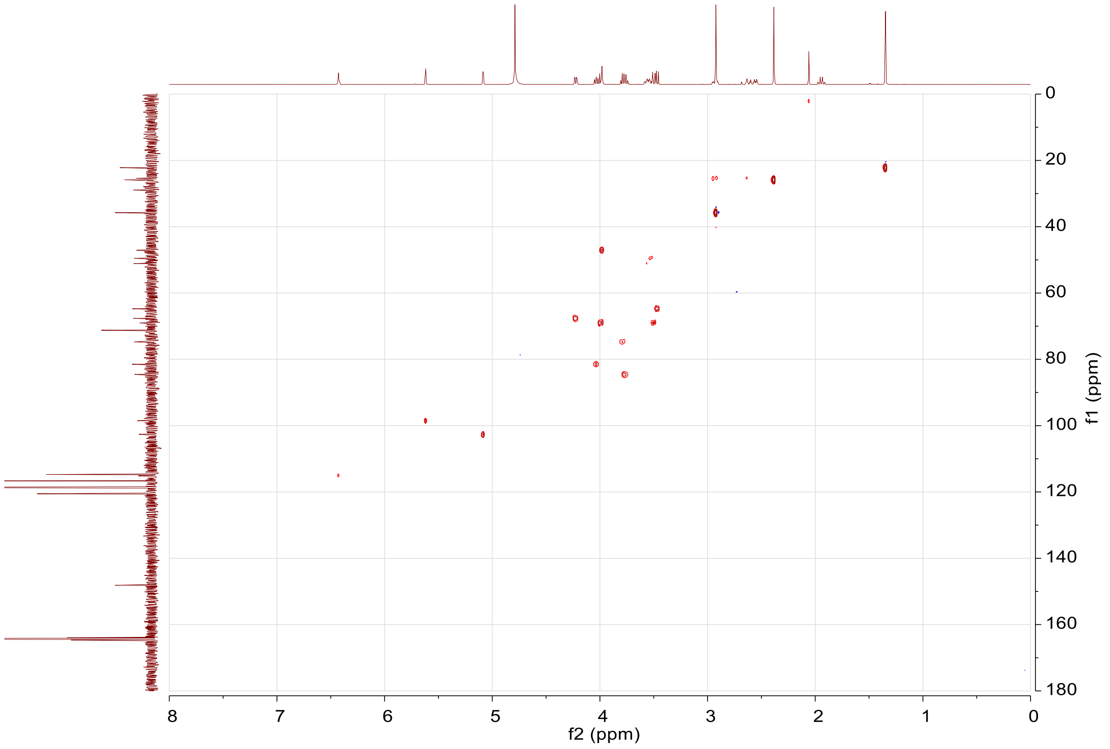

**e**, HMBC spectrum (600 MHz) of oxo-verdamicin (**6**).

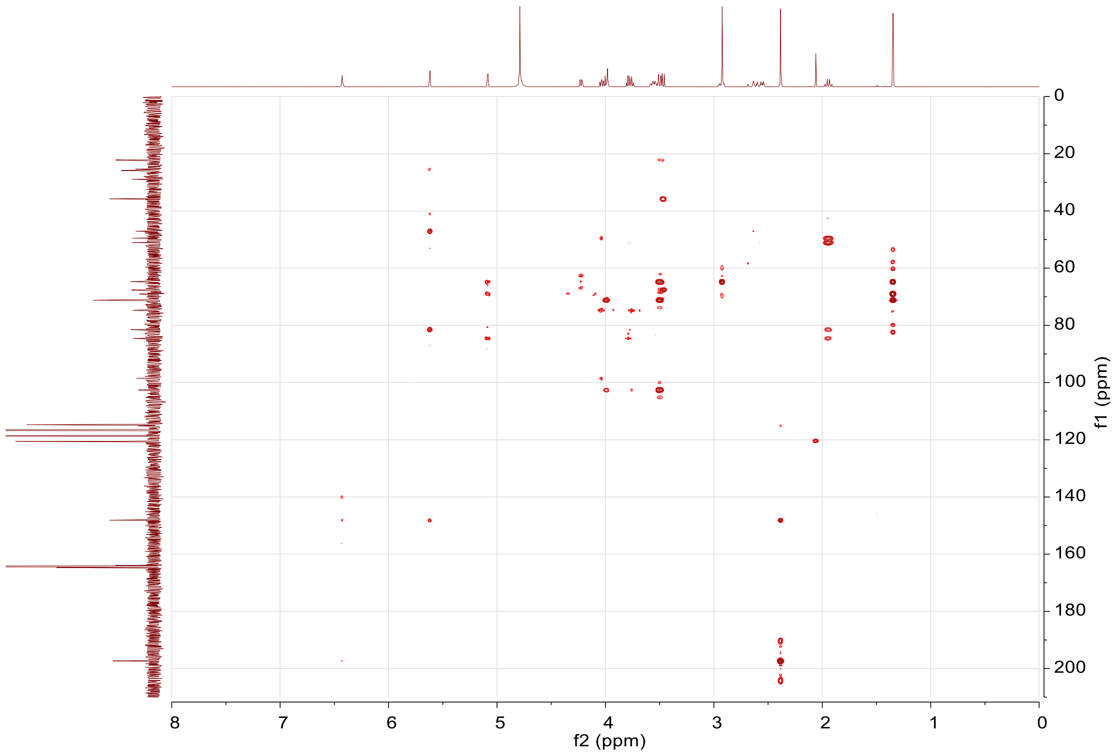


**f**, NOESY spectrum (600 MHz) of oxo-verdamicin (**6**).

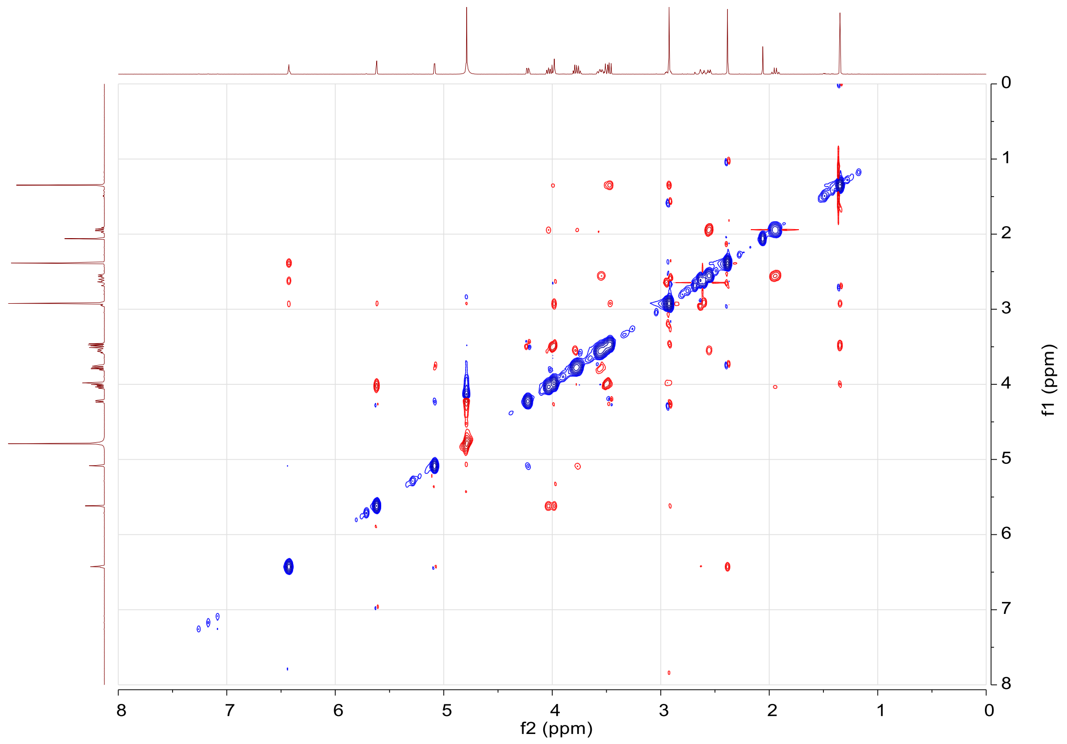

**g**, Structure and conformation determination of oxo-verdamicin (**6**).

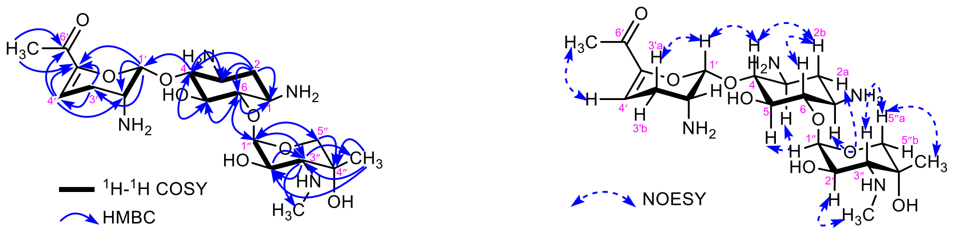


oxo-verdamicin (**6**) oxo-verdamicin (**6**)

**h.** $^{1}$H NMR spectrum (400 MHz, $D_2O$) analysis of oxo-verdamicin (**6**) and sisomicin (**7**) in acidic and basic conditions.

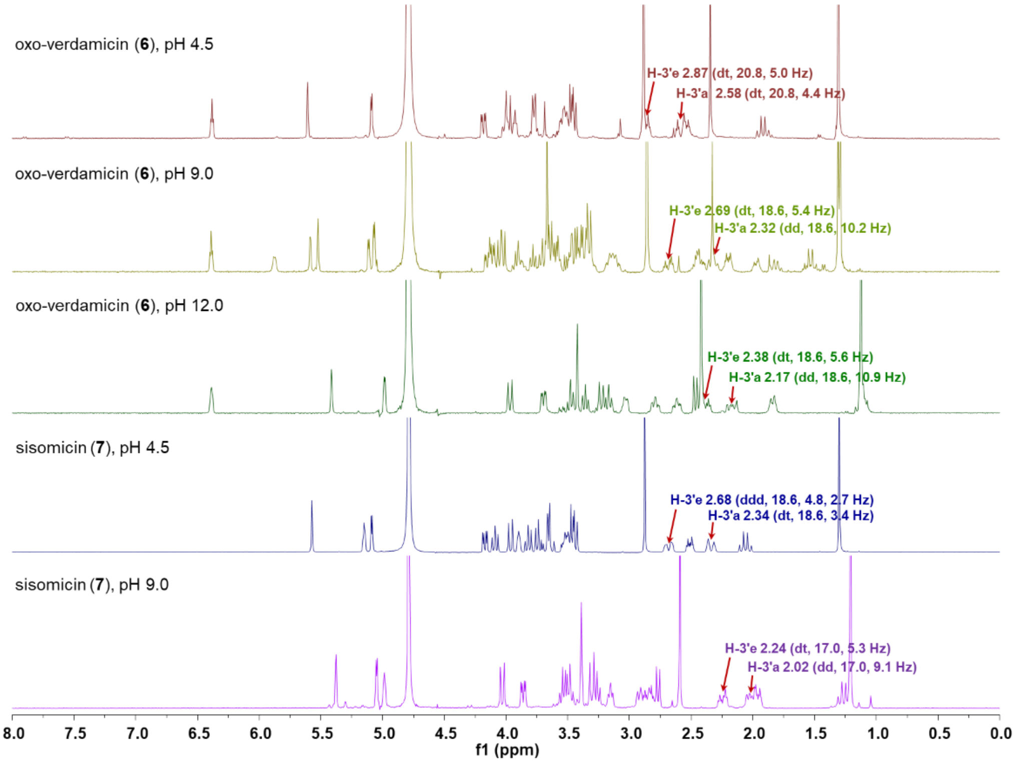


**i.** Conformational changes of oxo-verda (**6**), sisomicin (**7**), verdamicin C2a (**8**) and verdamicin C2 (**9**) in basic and acidic conditions.

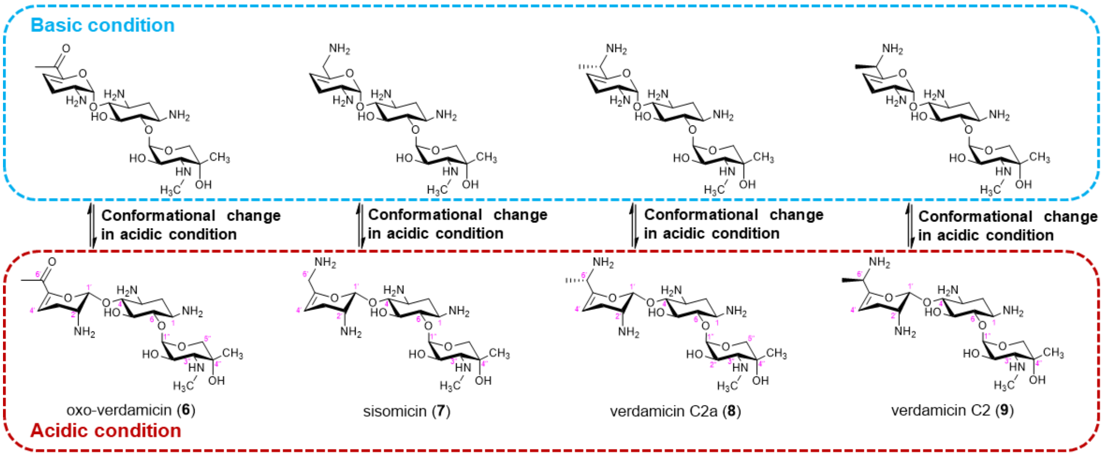

**Supplementary Fig. 4 | Confirmation of gene complementation mutants.**

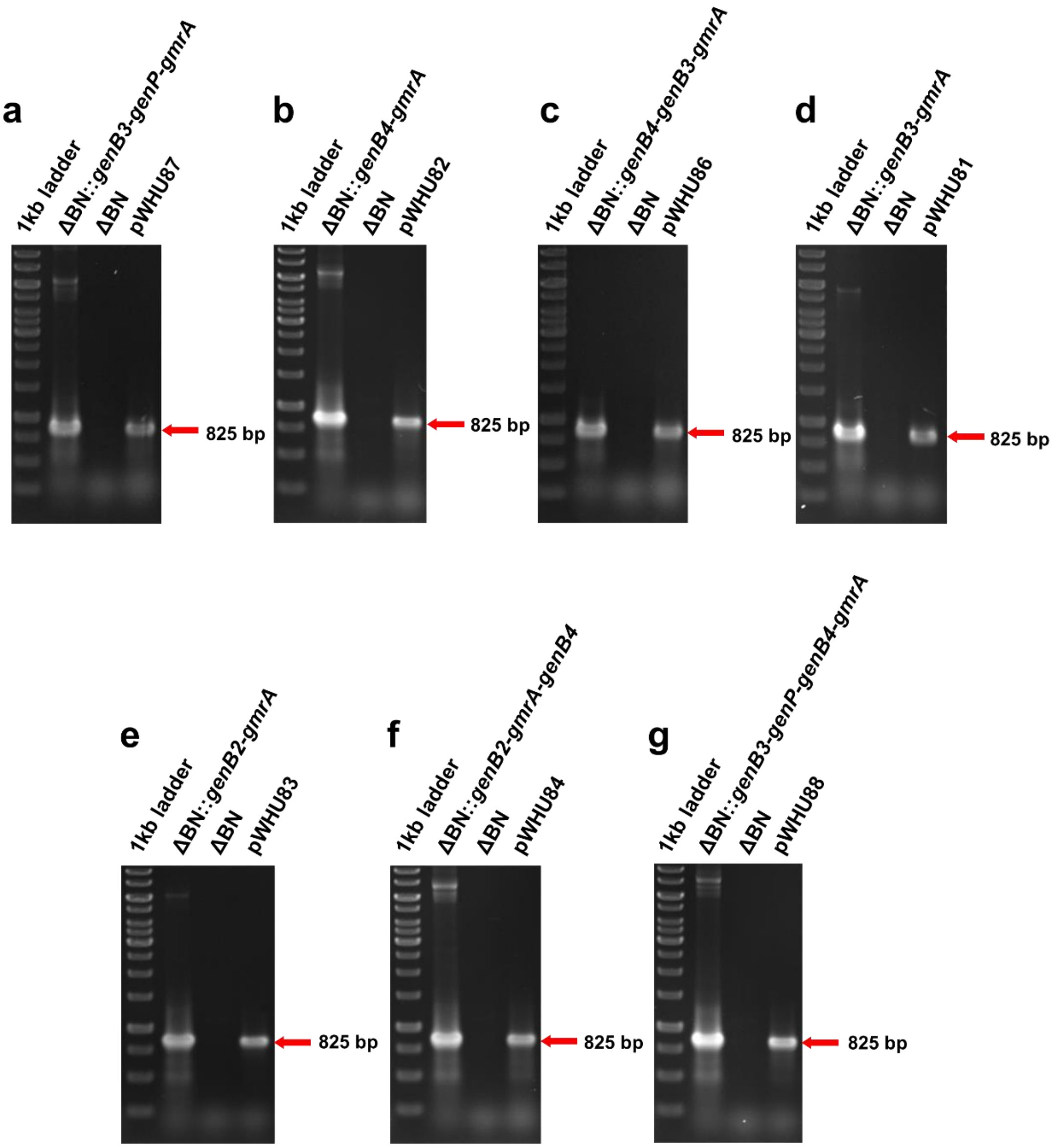


Check primers used for checking *gmr* are EP-gmrA-CK1 and EP-gmrA-CK2 listed in Supplementary Table 4. The red arrows indicate the expected size of PCR fragments of the complemented gene *gmrA*.

**Supplementary Fig. 5 | LC-ESI-HRMS analysis of products in feeding experiments.**

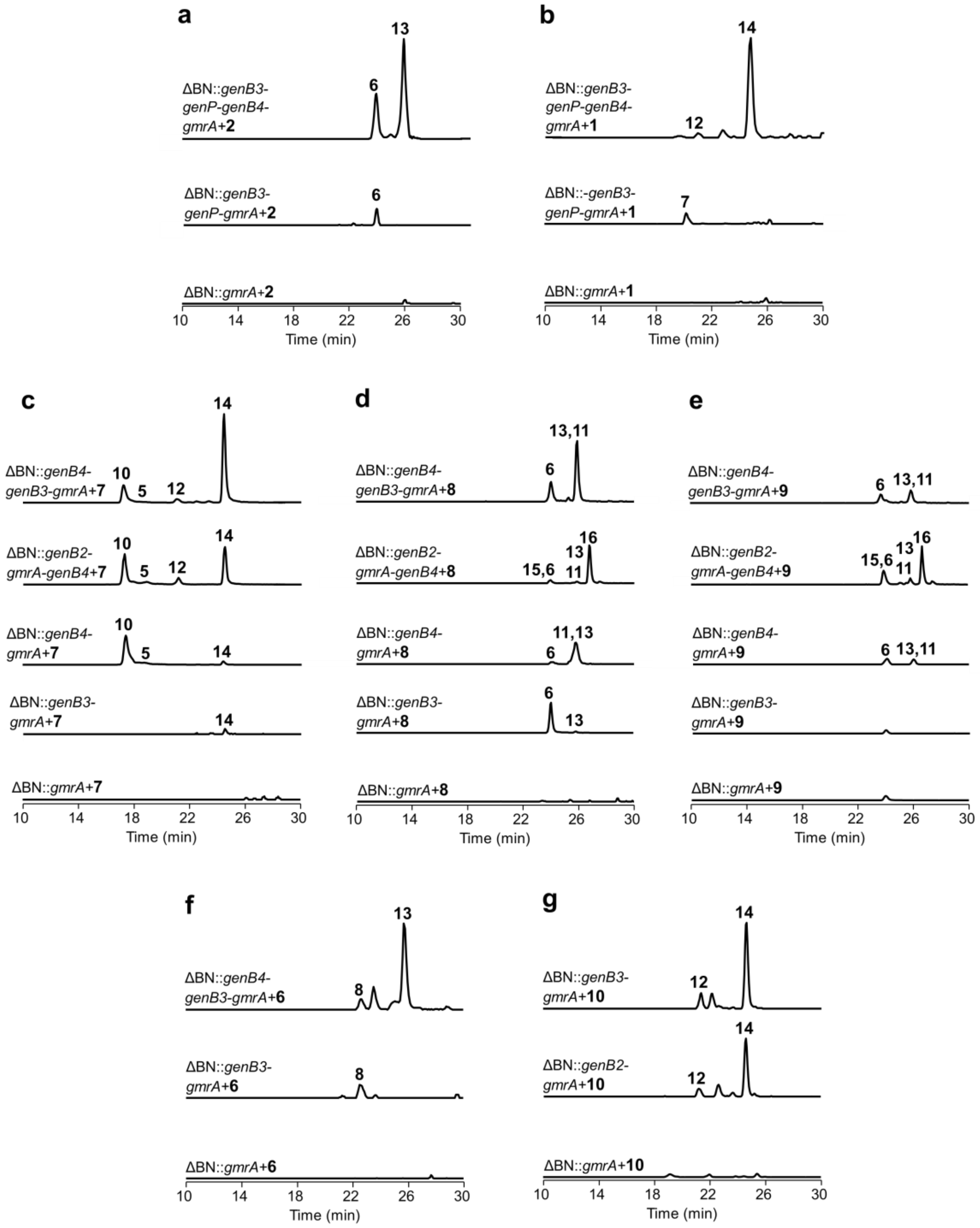


Extracted ion chromatogram trance of products in feeding experiments using **a**, JI-20Ba (**2**); **b**, JI-20A (**1**); **c**, sisomicin (**7**); **d**, verdamicin C2a (**8**); **e**, verdamicin C2 (**9**); **f**, oxo-verdamicin (**6**); and **g**, oxo-C1a (**10**) as substrates to ΔBN-based complementary mutants.

**Supplementary Fig. 6 | NMR analysis of gentamicin intermediates verdamicin C2a (8) and verdamicin C2 (9) in $D_2O$.**

**a**, $^1H$ spectrum (600 MHz) of verdamicin C2a (**8**).

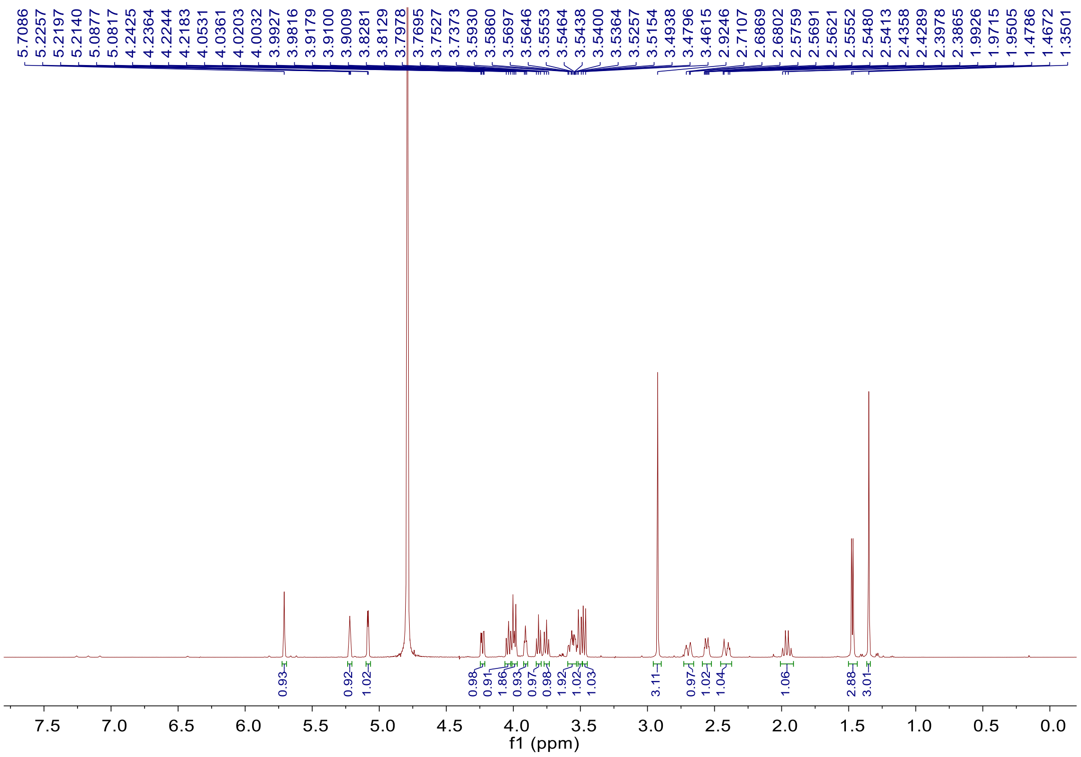


**b**, $^1H$-$^1H$ COSY spectrum (600 MHz) of verdamicin C2a (**8**).

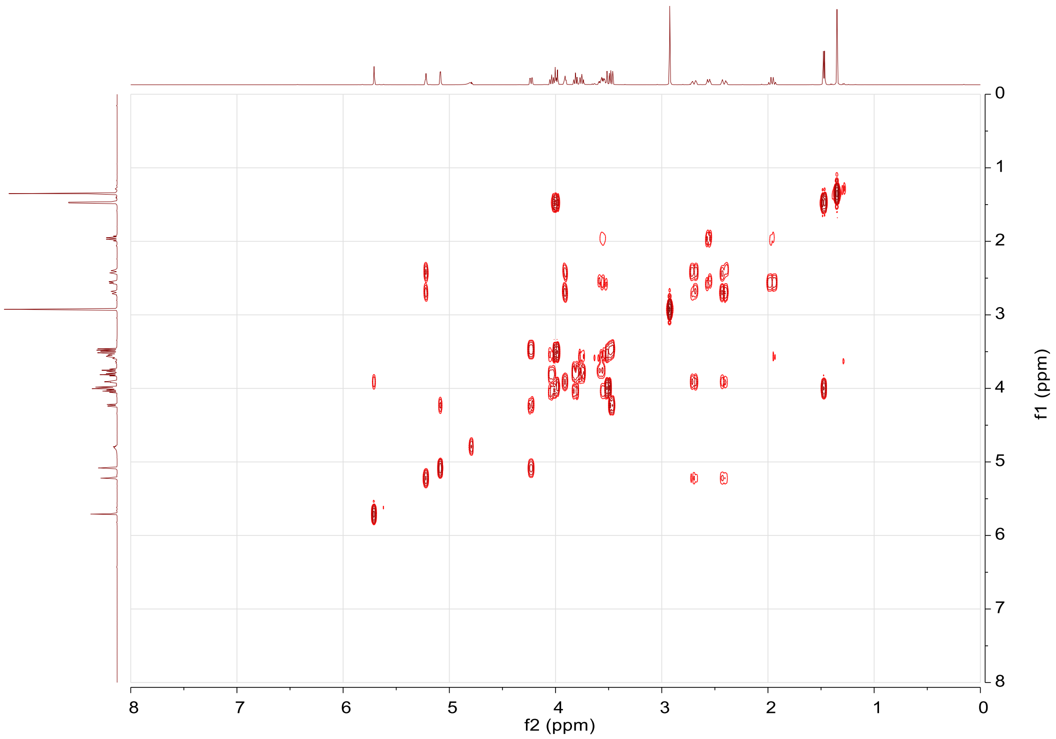

**c**, NOESY spectrum (600 MHz) of verdamicin C2a (**8**).

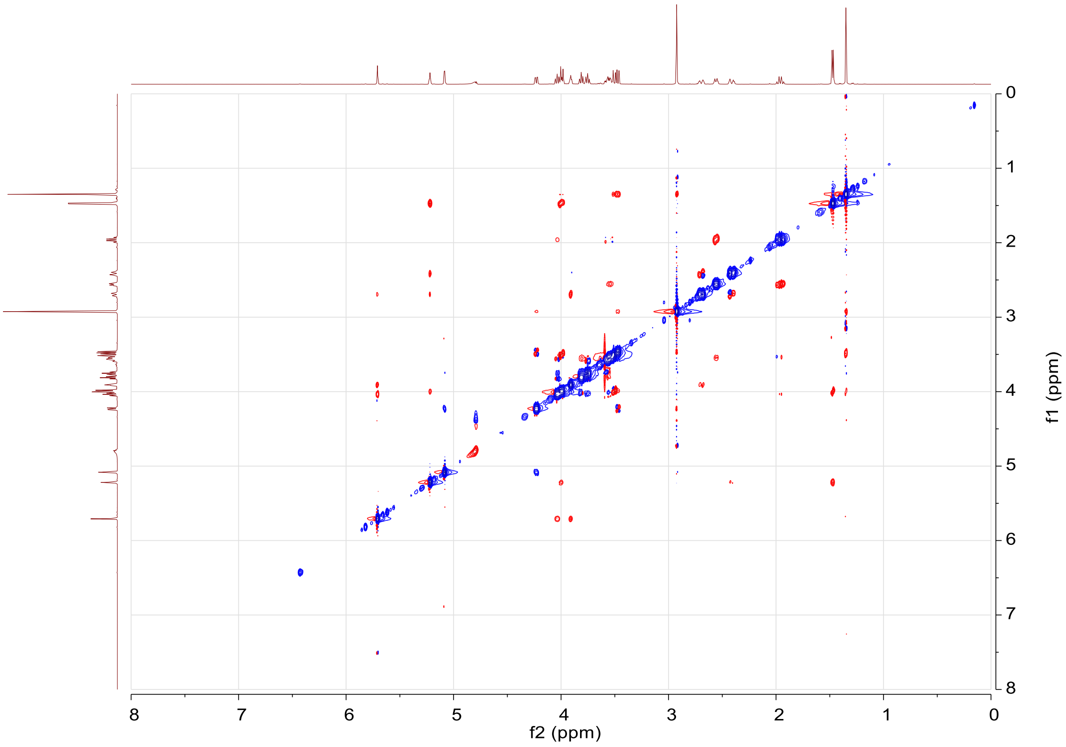


**d**, [1]H spectrum (600 MHz) of verdamicin C2 (**9**).

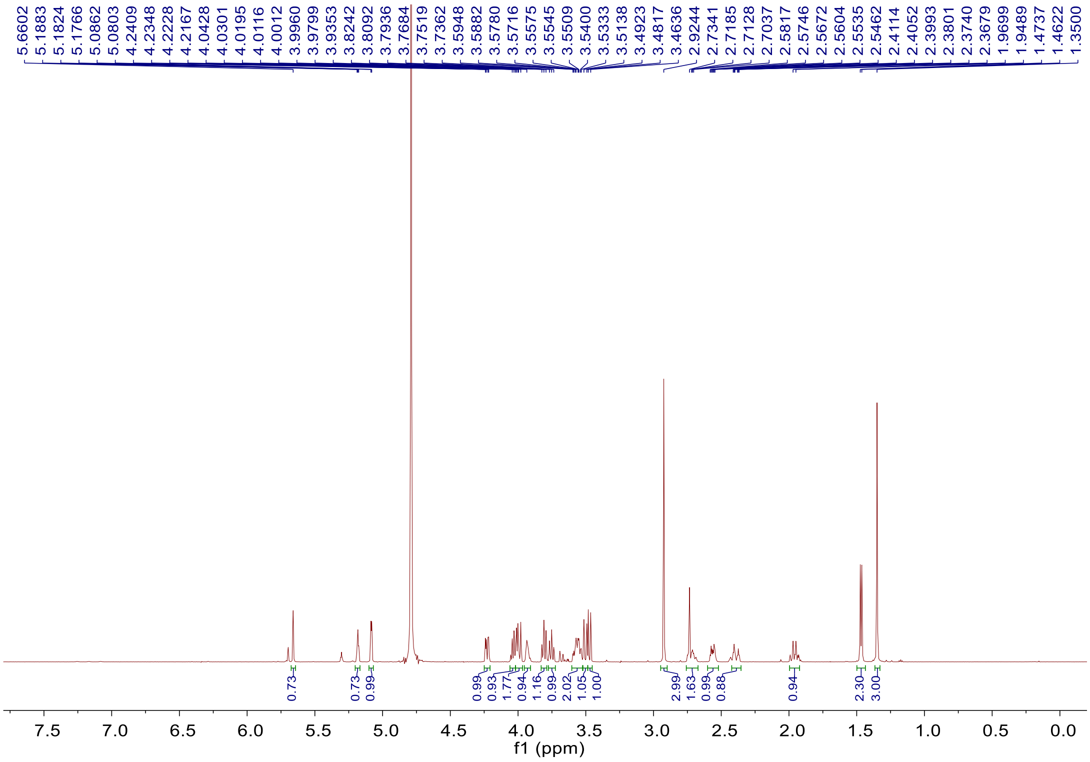

**e**, $^{1}H$-$^{1}H$ COSY spectrum (600 MHz) of verdamicin C2 (**9**).

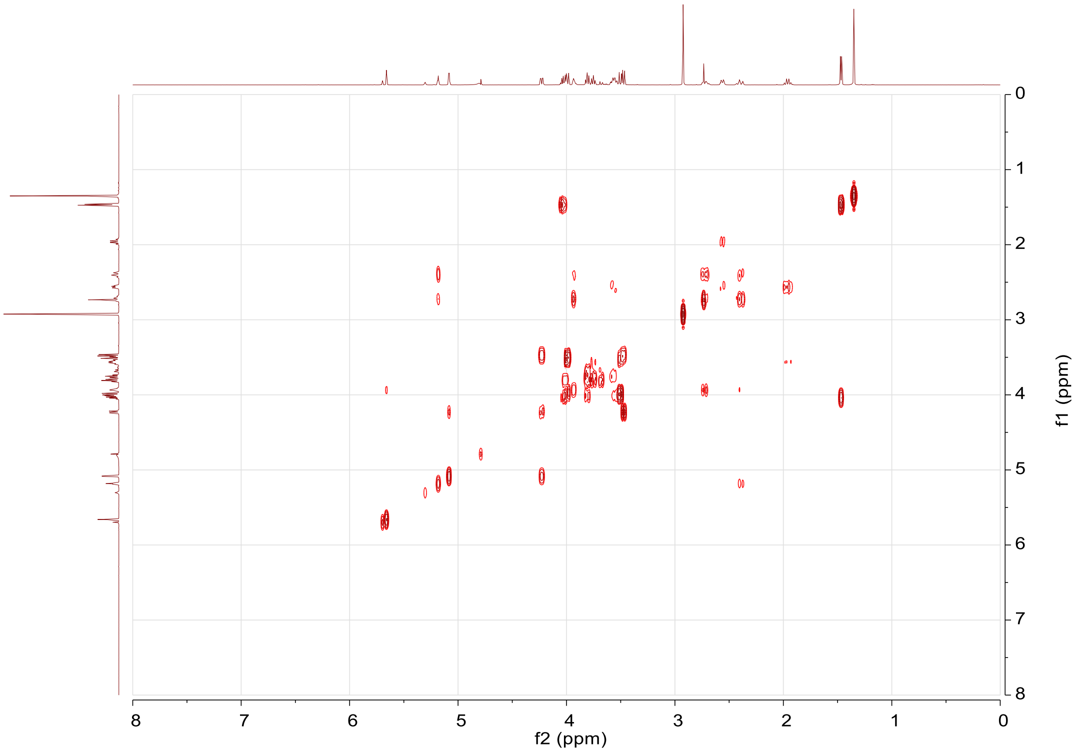


**f**, NOESY spectrum (600 MHz) of verdamicin C2 (**9**).

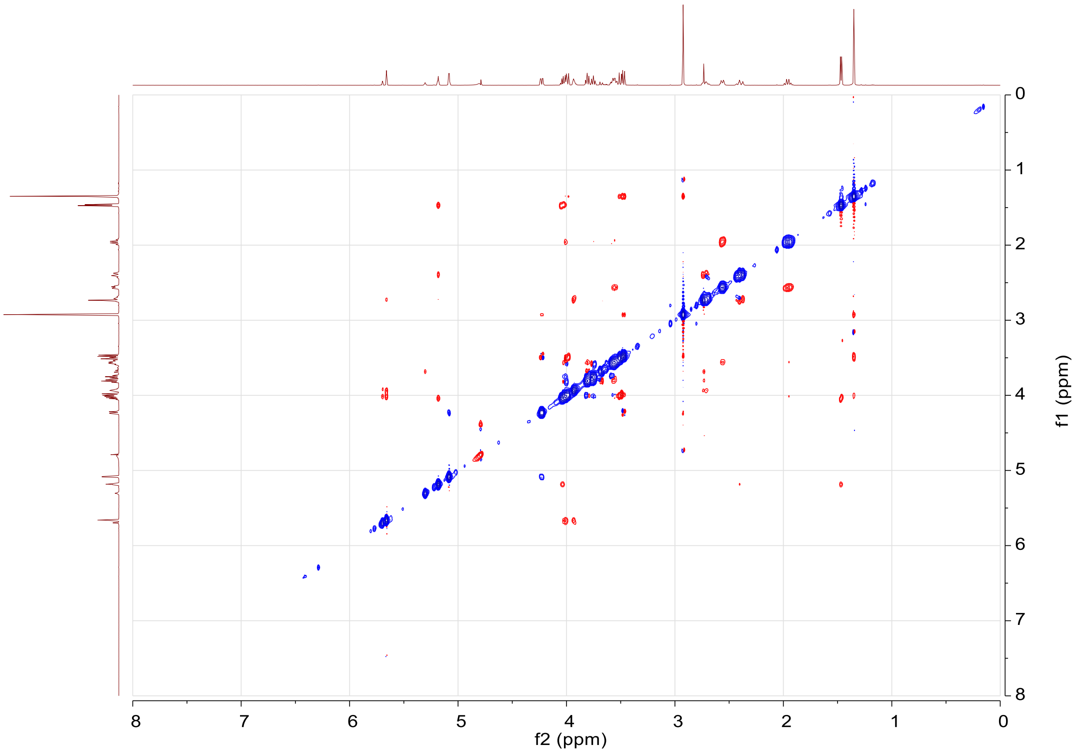

**g**, Structure and conformation determination of verdamicin C2a (**8**) and verdamicin C2 (**9**).

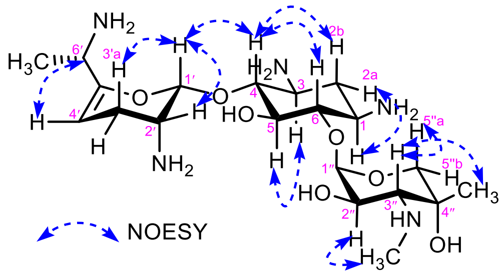

verdamicin C2a (**8**)

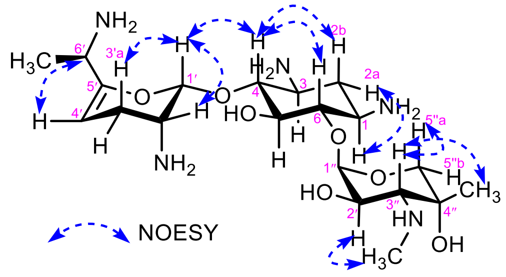

verdamicin C2 (**9**)

**Supplementary Fig. 7 | Verification of configuration of verdamicin C2a (8) and verdamicin C2 (9) by LCMS comparison with authentic synthetic standards of known stereochemistry[6].**

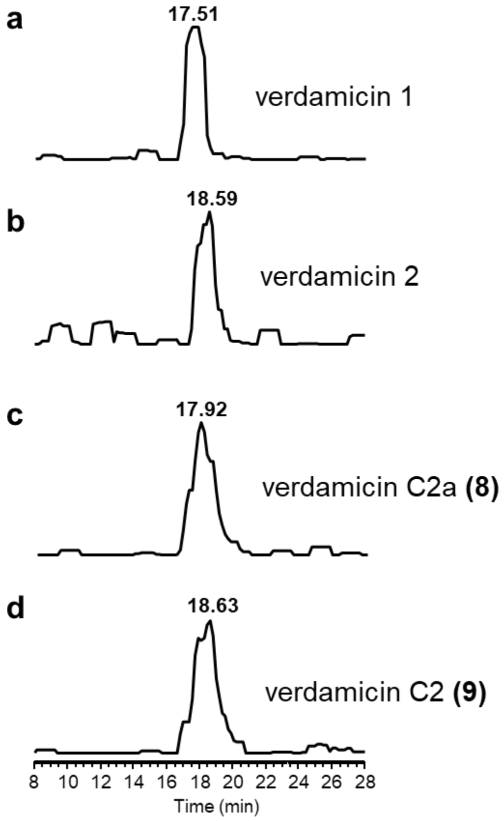


LC-ESI-MS extracted ion chromatograms for *m/z* 484.3 $[M+Na]^+$ are shown. **a**, the verdamicin isomer (verdamicin 1) produced by GenB3-catalysed transamination of oxo-verdamicin (**6**); **b**, the other verdamicin epimer (verdamicin 2) isolated from *in vivo* experiments; **c**, synthetic verdamicin C2a (**8**); **d**, synthetic verdamicin C2 (**9**). Note: **a** and **b** were run on different days to **c** and **d** and on this LC-MS system there was some day-to-day variation of retention times. However verdamicin1 always eluted before verdamicin 2 and verdamicin C2a always eluted before verdamicin C2.

**Supplementary Fig. 8 | LC-ESI-HRMS analysis of the influence of C-6' stereochemistry on didehydroxylation in vitro.**

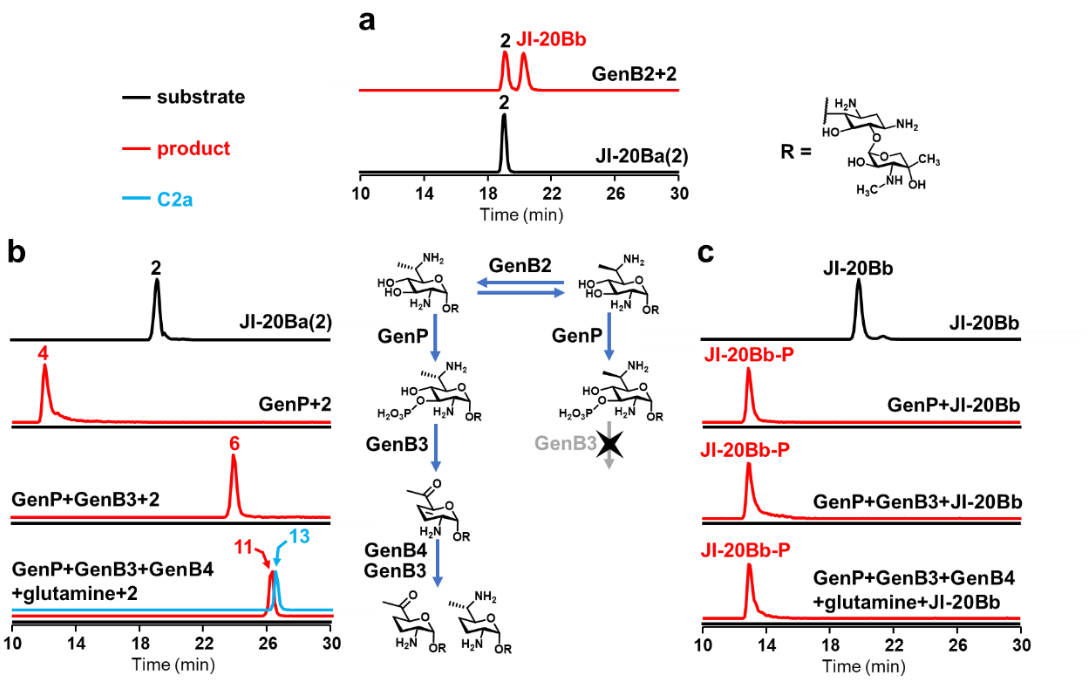


**a**, JI-20Ba was epimerized to its 6'-epimer JI-20Bb by GenB2; **b**, JI-20Ba and **c**, JI-20Bb were compared as the substrates of GenP, GenB3 and GenB4. Black and red lines refer to extracted ion chromatograms of the substrate and products, respectively. C2a (**13**) in assay products is particularly indicated by blue lines due to the overlapping of its retention time with that of oxo-C2a (**11**).

**Supplementary Fig. 9 | Superposition of PLP-dependent enzymes from gentamicin biosynthesis.**

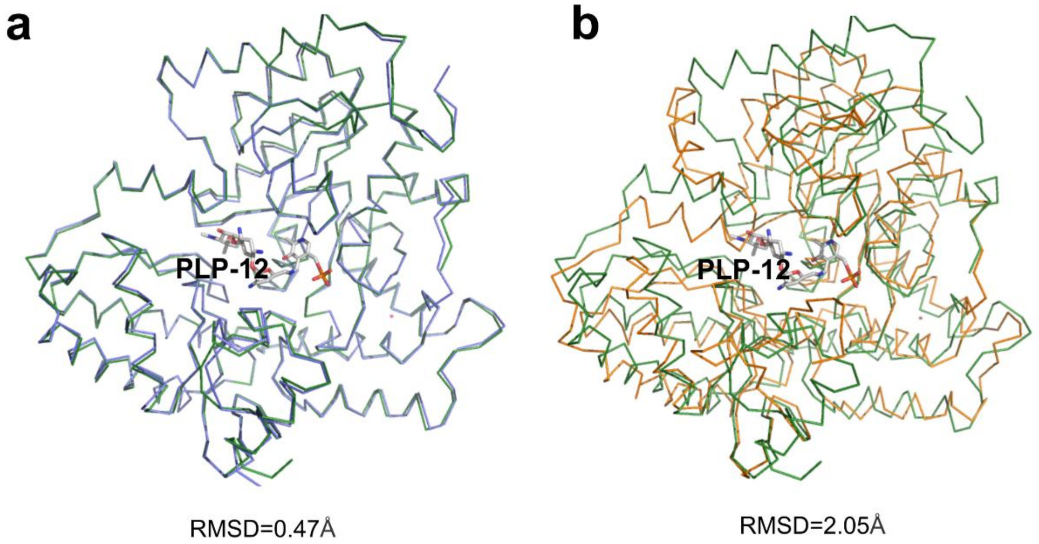


**a**, GenB3-PLP (blue) and GenB4-PLP-**12** (green); **b**, GenB4-PLP-**12** (green) and GenB1-PLP (orange).

**Supplementary Fig. 10 | Amino acid alignments for the PLP-dependent enzymes from gentamicin biosynthesis.**

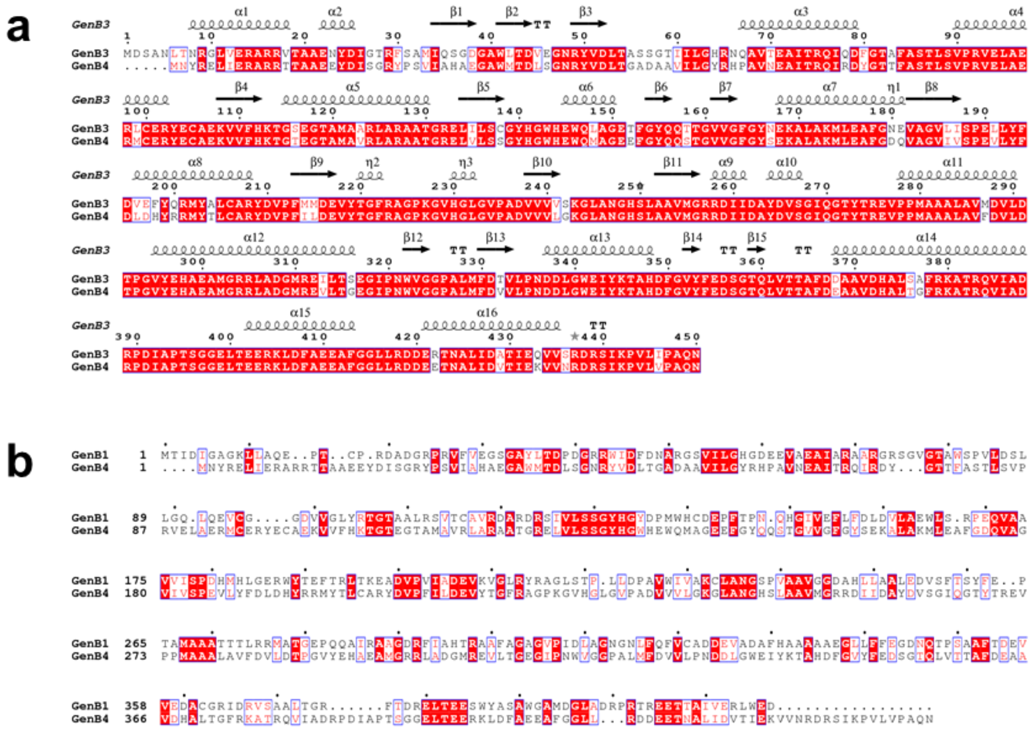

**a**, GenB3 and GenB4; **b**, GenB1 and GenB4.

**Supplementary Fig 11. Schiff base of GenB4 and GenB3. Internal aldimine of Lys238 and Lys243 with PLP electron density map contours for a, GenB4 and b, GenB3.**

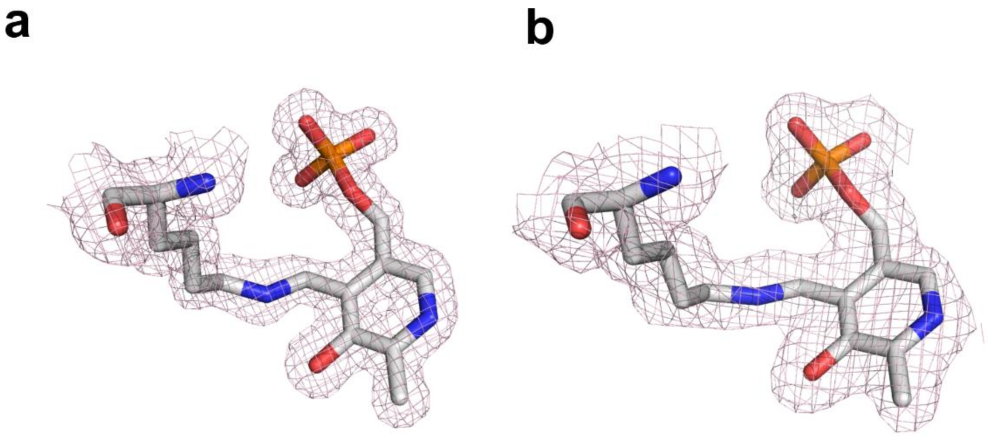


**Supplementary Fig. 12 | Superposition of GenB4-PLP (blue) and GenB4-PLP-12 (orange).**

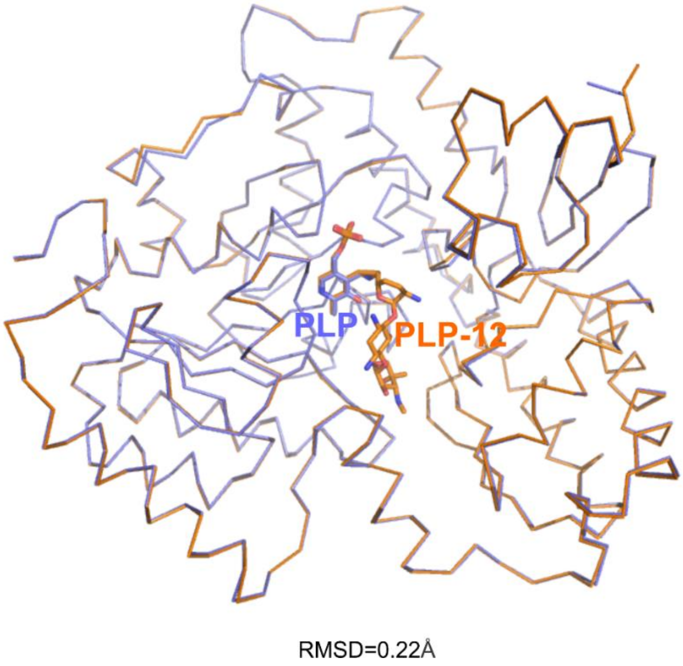

**Supplementary Fig. 13 | *Gem*-diamino observed in the protomer A of GenB4-PLP-12.**

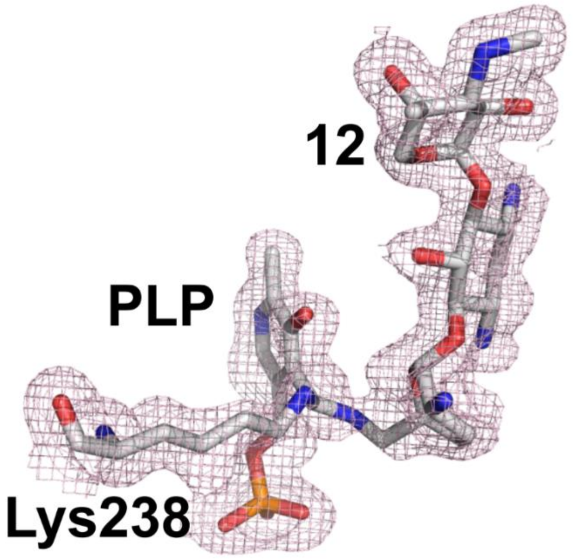


**Supplementary Fig. 14 | Superposition of active sites of a, GenB4-PLP (green) and GenB4-PLP-12 (beige) and b, GenB4-PLP-12 (protomer A, beige) and GenB4-PLP-12 (protomer B, gray).**

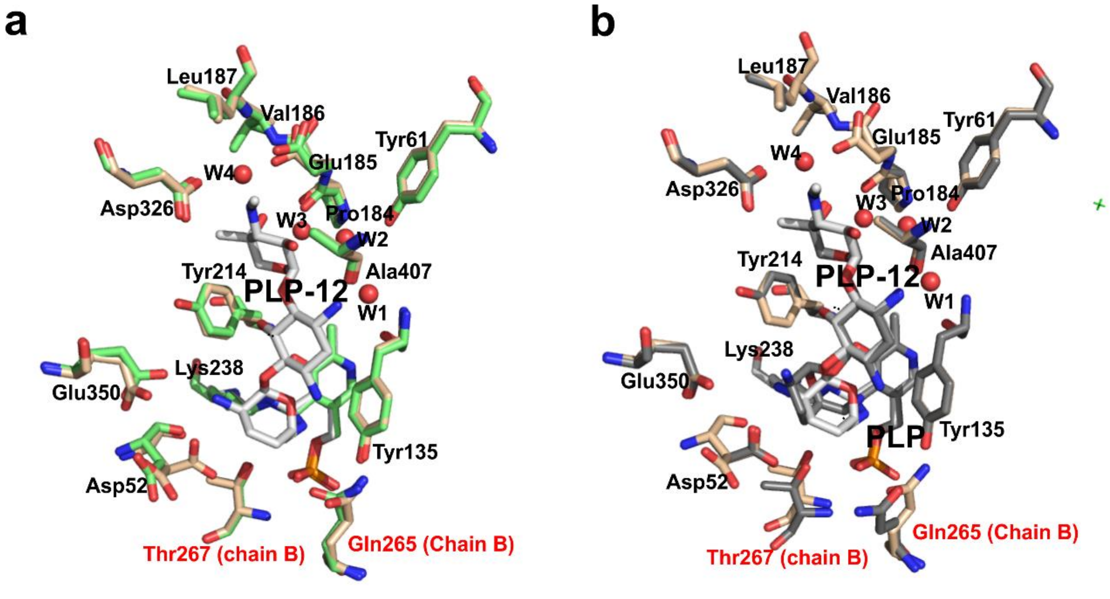

**Supplementary Fig. 15 | Potential electrostatic surface of GenB4 (a) and GenB3 (b). A close-up of each active site is indicated by a box.**

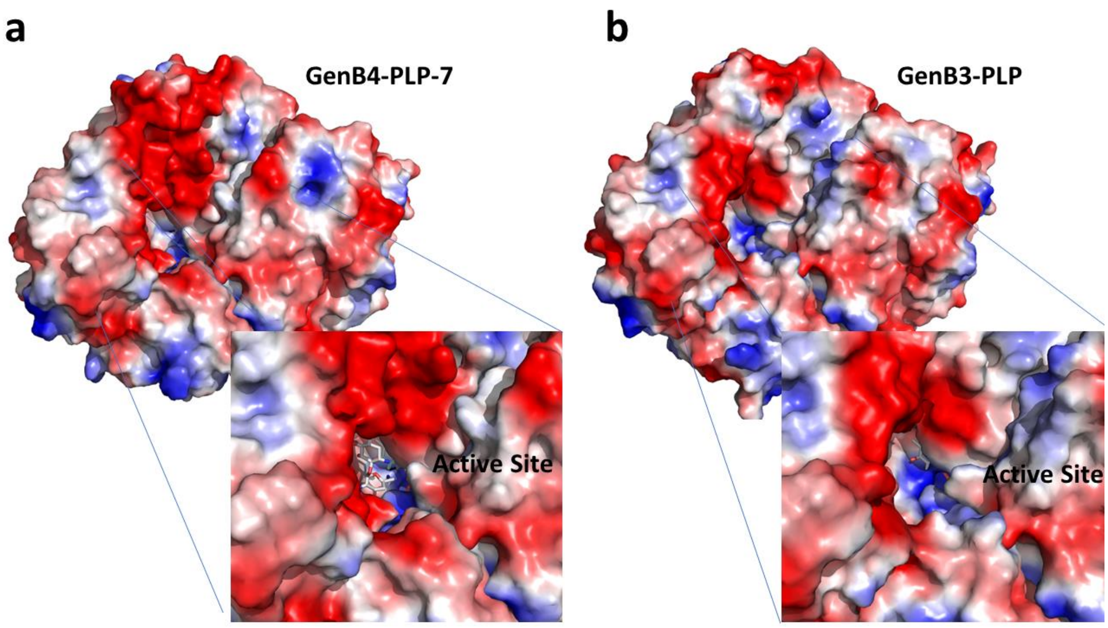
